# ATF4 activates a transcriptional program that chronically suppresses mTOR activity promoting neurodegeneration in Parkinson’s disease models

**DOI:** 10.1101/2025.06.09.658667

**Authors:** Matthew D. Demmings, Erica A. Kane, Elizabeth C. Tennyson, Kate Hurley, Joy Zhao, Nicholas A. Cruickshanks, Victoria Ciz, Jordan M. Krupa, Stephen H. Pasternak, Sean P. Cregan

**Author notes:** Corresponding Author: Sean P. Cregan.

## Abstract

The Integrated Stress Response (ISR) is a cell signaling pathway that enables cells to adapt to diverse cellular stresses. Conversely, during chronic/unmitigated cellular stress the ISR becomes maladaptive and has been implicated in a range of neurodegenerative conditions including Parkinson’s Disease (PD). However, the mechanisms by which maladaptive ISR/ATF4 signaling contributes to neurodegeneration have not been elucidated. In this study we establish a critical mechanism by which chronic ISR activation becomes maladaptive and promotes neurodegeneration in neurotoxin and α- synucleinopathy models of PD *in vitro* and *in vivo*.

Specifically, we demonstrate that chronic activation of ATF4, the central transcription factor of the ISR, promotes neurodegeneration by regulating the transcriptional induction of SESN2, DDIT4 and Trib3 that co-operate to suppress both mTORC1 and mTORC2 activity. Furthermore, we demonstrate that ATF4-mediated suppression of mTORC1/2 activity promotes dopaminergic neuronal death in PD models by facilitating the activation of the pro-apoptotic BCL-2 family protein PUMA. Taken together, we have discovered a novel maladaptive ISR/ATF4 signaling pathway leading to chronic suppression of mTORC1/2 activity resulting in PUMA-mediated neuronal death that may have therapeutic implications in a range of neurodegenerative conditions that exhibit chronic ISR activation.

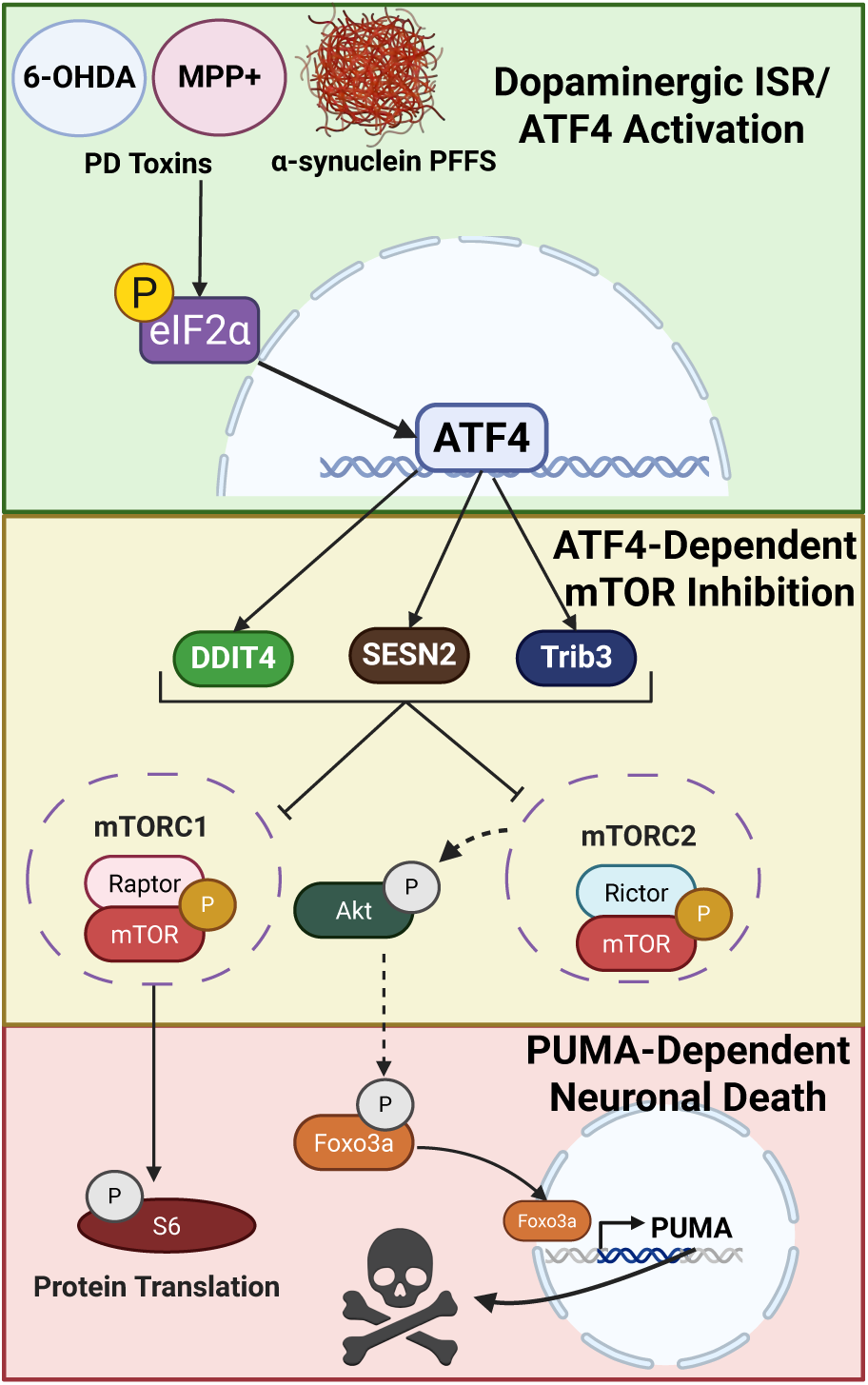

## Introduction

Parkinson’s disease (PD) is a neurodegenerative disease in which dopaminergic neurons of the substantia nigra progressively degenerate resulting in the loss of striatal dopamine and characteristic motor symptoms including bradykinesia, resting tremor and gait impairment (*1*). The accumulation of intracellular aggregates containing α-synuclein in Lewy bodies or Lewy neurites is a definitive hallmark of PD (*2*). Current treatments temporarily mitigate some of the clinical symptoms but do not address the underlying neurodegenerative processes. Therefore, there is a critical need to identify the cellular and molecular processes driving neurodegeneration in PD to guide the development of disease modifying therapeutics.

Numerous studies point to the central roles of proteostasis dysfunction, mitochondrial dysfunction, and oxidative stress in dopaminergic (DA) neuron degeneration in PD. While the majority of PD cases are idiopathic, familial-linked PD-related genes provide insight into mechanisms potentially contributing to disease. Causal mutations in the PD-related gene SNCA that codes for α-synuclein have been shown to produce aggregation prone forms of the protein and is associated with early onset disease (*3*, *4*). Experimental production of α-synuclein aggregates in transgenic mice over-expressing mutant human α-synuclein or in mice seeded with α-synuclein preformed fibrils has been shown to drive neurodegeneration (*5–7*). Furthermore, mutations in the PD-associated genes GBA1 and LRRK2 have been shown to compromise lysosome and endolysosome function leading to deficits in autophagic clearance of α-synuclein aggregates (*8–11*). On the other hand, numerous studies support a causal role of mitochondrial dysfunction and oxidative stress in PD associated neurodegeneration. The PD-genes PARKN, PINK1 and DJ-1 have been implicated in regulating oxidative stress and mutations in these genes have been associated with deficits in mitophagy resulting in the accumulation of dysfunctional mitochondria (*12*–*16*). Furthermore, neurotoxins such as MPTP, 6-OHDA, rotenone, and paraquat induce selective dopaminergic neuron degeneration in animals by causing mitochondrial dysfunction and oxidative stress (*17–20*). Moreover, it was shown that selective disruption of mitochondrial complex-I function in dopaminergic neurons is sufficient to induce progressive parkinsonism in mice (*21*). A prevailing hypothesis is that dopaminergic neurons are intrinsically sensitive to mitochondrial dysfunction due to their high bioenergetic activity associated with their extensive neurite arborization and tonic activity (*22*, *23*).

The Integrated Stress Response (ISR) is a cell signaling pathway that is activated in response to diverse stress stimuli including oxidative stress and proteostatic stress prevalent in neurodegenerative conditions (*24*). Indeed, chronic ISR activation has been implicated in a wide range of brain disorders including Parkinson’s disease (*25–27*). The primary function of the ISR is to attenuate general protein translation and enable the selective synthesis of stress responsive proteins to restore cell homeostasis, however during prolonged or unmitigated cellular stress the ISR becomes maladaptive and can promote cell death (*24*). Activating transcription factor-4 (ATF4) is the central transcription factor activated in the ISR and is known to induce the expression of genes involved in amino acid metabolism, redox homeostasis and protein folding that promote cell recovery (*28*). However, during prolonged ISR activation, ATF4 can induce the expression of genes associated with cell death (*29*–*31*). Elevated levels of ATF4 have been reported in cellular and animal models of PD as well as in *post-mortem* tissue of PD patients (*32*, *33*). Previously, we demonstrated that ATF4 expression is persistently upregulated and promotes dopaminergic neuronal death in neurotoxin and α-synucleinopathy models of PD (*34*). However, the mechanism by which chronic ISR/ATF4 activation becomes maladaptive and promotes neurodegeneration remains unclear and is the focus of this study.

Mammalian Target of Rapamycin (mTOR) is a serine/threonine protein kinase that integrates nutrient, growth factors and cellular energy inputs to regulate protein synthesis, metabolism and survival (*35*). mTOR forms two distinct complexes, mTORC1 and mTORC2, that respond to different stimuli and target distinct substrates. In the mTORC1 complex, mTOR associates with RAPTOR and phosphorylates several downstream targets including ribosomal S6K1 and 4E-BP1 to regulate protein synthesis and ULK1 to inhibit autophagy (*36*–*38*). In the mTORC2 complex, mTOR associates with RICTOR and phosphorylates AKT, PKCα and SGK1 to regulate survival, metabolism and cytoskeletal dynamics (*39*–*41*). In the nervous system, mTOR has been demonstrated to play an important role in long lasting forms of LTP and LTD associated with synaptic plasticity and learning and memory (*42–48*). Similarly, both mTORC1 and mTORC2 signaling have been implicated in the regulation of dopaminergic neural transmission, plasticity and dopamine-dependent behaviours (*49*–*51*). Furthermore, the survival promoting function of the mTORC2 target, AKT, in neurons has been widely documented (*52*, *53*). In recent years, there has been accumulating evidence demonstrating that mTOR signaling is dysregulated in PD models and during progression of PD in humans (*54*).

Previously, we demonstrated that ATF4 promotes neuronal death in neurotoxin and α- synucleinopathy models of PD in dopaminergic neuronal cultures (*34*). In this study we extend these findings and demonstrate that the ATF4 orthologue *Atfs-1* promotes dopaminergic neuronal dysfunction and degeneration in neurotoxin and α-synuclein PD models in *C. elegans in vivo*. Furthermore, we establish a critical mechanism by which chronic ISR/ATF4 activation becomes maladaptive and promotes neurodegeneration in PD models. Specifically, we demonstrate that chronic ISR/ATF4 activation promotes degeneration of dopaminergic neurons by suppressing mTORC1 and mTORC2 activity via the transcriptional induction of the mTOR inhibiting proteins SESN2, DDIT4 and Trib3. Furthermore, we demonstrate that ATF4-mediated suppression of mTOR activity promotes dopaminergic neuronal death in PD models by facilitating the activation of the pro-apoptotic BCL-2 family protein PUMA. Taken together, we have established a key maladaptive ISR/ATF4 signaling pathway that promotes neurodegeneration through suppression of mTORC1/2 activity that may have therapeutic implications in a range of neurodegenerative conditions that exhibit chronic ISR activation.

## Results

### Atfs-1-Loss of function in C. elegans reduces DA neuron dysfunction and neurodegeneration and enhances lifespan in PD neurotoxin and α-synuclein models

Having previously demonstrated that sustained activation of the ISR transcription factor ATF4 promotes dopaminergic (DA) neuron death in cellular models of PD (*34*), we investigated whether chronic ATF4 activation induces dopaminergic neuron dysfunction and degeneration *in vivo*. The nematode *Caenorhabditis elegans* (*C. elegans*) has become a valuable organism for modeling and investigating molecular mechanisms of neurodegenerative diseases including Parkinson’s disease (*55*). The essential components of synaptic transmission such as neurotransmitters, transporters, receptors, and ion channels are conserved in *C. elegans* (*56*). However, the nervous system of *C. elegans* is comparatively simple consisting of only 302 neurons in the adult animal, of which only 8 are dopaminergic (*57*). Furthermore, the short (∼20-30 day) lifespan and transparent anatomy of *C. elegans* enables assessment of neurodegeneration using neuron-specific fluorescent reporter transgenes over the animals’ lifespan (*58*–*60*). The *C. elegans* basic leucine zipper transcription factor ATFS-1 is the central regulator of the mitochondrial unfolded protein response (UPR^MT^) that is activated in response to mitochondrial dysfunction and the accumulation of unfolded proteins within the mitochondria (*61*). Similar to its mammalian ortholog ATF4, ATFS-1 is activated by mitochondrial dysfunction and translocates to the nucleus where it regulates the expression of genes that code for proteins involved in protein misfolding, metabolism and stress resistance in an effort to restore mitochondrial homeostasis (*62*). Previous studies have demonstrated that the UPR^MT^ is activated in *C. elegans* exposed to the PD neurotoxins MPP+ and 6-OHDA as well as in transgenic animals expressing α-synuclein (*63*). Therefore, to investigate the role of ATFS-1 in dopaminergic neurodegeneration in *C. elegans* PD models we compared the response of N2 wildtype animals and *C. elegans* expressing a loss-of-function mutant atfs-1 allele (*atfs-1(gk3094*)) hereafter referred to as *atfs-1 LoF*. To visualize dopaminergic neurons we crossed the *atfs-1 LoF* animals with a transgenic *C. elegans* line that constitutively expresses GFP specifically in dopaminergic neurons (*dat-1::GFP*). *C. elegans* hermaphrodites possess 4 cephalic (CEP), two anterior deirid (ADE) and two posterior deirid (PDE) dopaminergic neurons (*57*). To assess DA neuron survival, we counted the number of intact GFP+ CEP- and ADE- dopaminergic neuron cell bodies in each animal as a function of time after treatment with MPP+ or 6-OHDA as previously described (*58*–*60*). As shown in figure 1A, DA neuron survival was significantly higher in *atfs-1 LoF* animals as compared to wildtype (N2) animals following treatment with either MPP+ (p<0.0001) or 6-OHDA (p<0.0001) suggesting that chronic ATFS- 1 activation by PD neurotoxins promotes DA neuronal death *in vivo*. Next, we assessed dopaminergic neuron function in wildtype and *atfs-1 LoF* animals by assessing a dopamine-regulated behavior referred to as the Basal Slowing Rate (BSR). The BSR assay provides a quantitative measure of mechanosensory detection of bacterial food resulting in dopamine-dependent slowing of locomotion (*64*). The absence of this slowing behavior is indicative of a dopamine deficit. We used an automated video tracking system to analyze locomotive behavior in animals and measured the locomotion ratio in the presence versus absence of a food source. Quantification of BSR over a 12-day period revealed that *atfs-1 LoF* animals retained a greater BSR than wildtype animals in the presence of MPP+ (p=0.0059) or 6-OHDA (p=0.0054) (Fig. 1B). We also investigated the effects of chronic ATFS-1 expression in a *C. elegans* model of α-synucleinopathy. Since *C. elegans* do not express endogenous α- synuclein we utilized a transgenic line that expresses human α-synuclein fused to YFP under control of a muscle specific promoter (*unc-54p::α-Syn-YFP*) that develops extensive intracellular α-synuclein aggregation (*65*). The α-Syn expressing animals were crossed with *atfs-1 LoF* animals or maintained on the wildtype (N2) background. Similar to the PD neurotoxin models, α-Syn expression attenuated BSR in wildtype (N2) animals to a significantly greater extent than in *atfs-1 LoF* animals (p=0.0021) (Fig. 1B). These results suggest that chronic activation of ATFS-1 promotes dopaminergic dysfunction in neurotoxin and α-synuclein PD models *in vivo*. We next performed lifetime survival assays to determine whether chronic ATFS-1 activation affects longevity in the PD models. We did not observe any difference in lifespan between untreated N2 and *atfs-1 LoF* animals (Fig. 1C). However, wildtype (N2) animals exhibited significantly reduced lifespan compared to *atfs-1 LOF* animals following treatment with either MPP+ (p=0.0178) or 6-OHDA (p=0.0053) (Fig. 1C).

**Figure 1.**
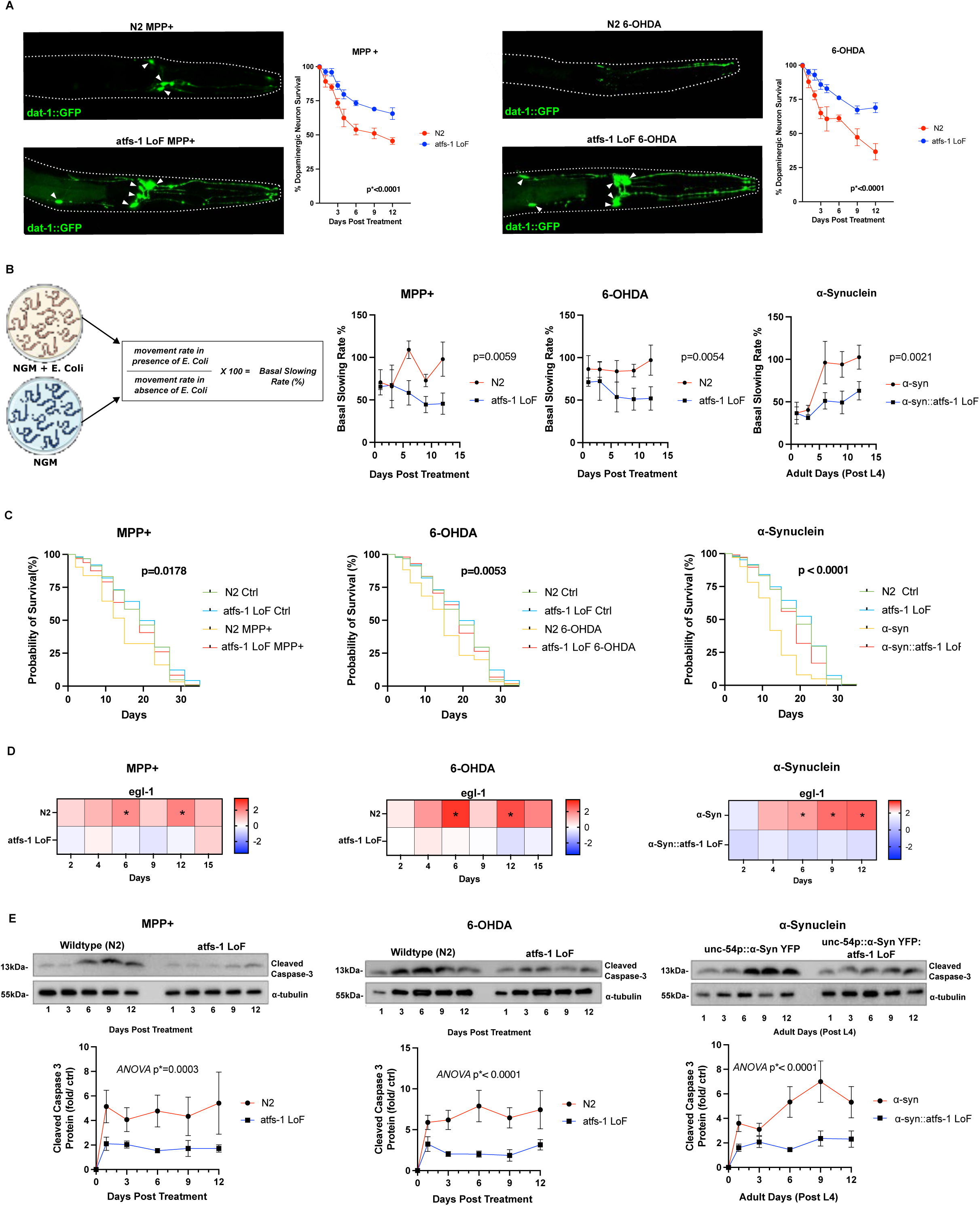
*Atfs-1* loss of function in *C. elegans* reduces DA neuron dysfunction and degeneration and enhances lifespan in neurotoxin and α-synuclein PD models. **A.** *dat-1:GFP::N2* and *dat-1:GFP::atfs-1 LoF C. elegans* were treated with MPP+ [100uM] or 6-OHDA [100uM] and dopaminergic neuron survival was scored at the indicated time points and reported as the mean +/- SEM (n=4, 2-way-ANOVA). Representative images of GFP labeled dopaminergic neurons (arrowheads). **B.** Basal Slowing Response (BSR) was determined for N2 and *atfs-1 LoF* animals treated with MPP+ [100uM] or 6-OHDA [100uM] and *a-syn* expressing animals or *a-syn::atfs-1 LoF* animals at the indicated times (n=5, 2-way-ANOVA). **C.** Kaplan-Meier survival curves of wildtype (N2) and *atfs-1 LoF C. elegans* treated with MPP+ [100uM] or 6-OHDA [100uM] and Wildtype (N2), *α-syn*, *atfs-1 LoF*, and *atfs-1 LoF::α-syn* animals. Statistical significance was determined by Log-rank (Mantel Cox) test. **D.** RNA was isolated from N2 animals or *atfs-1 LoF* animals treated with MPP+ [100uM] or 6-OHDA [100uM] and *α-syn* or *α-syn::atfs-1LoF* animals at indicated time points. Heat maps represent fold change in egl-1 mRNA levels as determined by qRT-PCR (n=3, 2-way-ANOVA, *p<0.05). **E.** Protein was isolated from N2 animals or *atfs-1 LoF* animals treated with MPP+ [100uM] or 6-OHDA [100uM] and *α-syn* or *α-syn::atfs-1LoF* animals at indicated time points. Cleaved caspase-3 protein levels were assessed by western blot analysis and quantified by densitometry and normalized to the beta-tubulin loading control (n=3, 2-way- ANOVA).

Similarly, α-synuclein overexpression reduced animal lifespan to a significantly greater extent in wildtype (N2) animals as compared to *atfs-1 LOF* animals (P<0.0001) (Fig. 1C). Consistent with this, PD neurotoxins and α-synuclein overexpression increased mRNA levels of the cell death effector egl-1 and induced caspase activation to a significantly greater extent in N2 animals as compared to *atfs-1 LOF* animals although these measures are not specific to dopaminergic neurons (Fig. 1D and 1E). Taken together these results suggest that chronic activation of ATFS-1 promotes dopaminergic neuronal dysfunction, neurodegeneration and decreases lifespan in neurotoxin and α-synucleinopathy models of PD in *C. elegans*.

### Transcriptome analysis identifies mTOR signaling as a key ATF4 regulated pathway in neurons exposed to oxidative/ proteostatic stress

We next sought to uncover the mechanisms by which sustained ATF4 activation mediates neuronal degeneration. We first conducted transcriptomic analysis of ATF4-dependent gene expression in neurons treated with arsenite, which is known to cause both oxidative and proteostatic stress that have been implicated in several neurodegenerative conditions including PD (*cc*, *c7*). Arsenite-treated cortical neurons exhibit a robust and sustained induction of ATF4 (Figure 2A). Furthermore, arsenite induced a significant increase in apoptotic neurons in wildtype but not ATF4-deficient neuronal cultures indicating that ATF4 is required to promote neuronal death (Figure 2B). Therefore, we performed microarray analysis using the Affymetrix GeneChip Platform on mRNA isolated from arsenite treated *ATF4+/+* and *ATF4-/-* neurons to identify ATF4-responsive genes and signaling pathways induced by oxidative/proteostatic stress. Differentially Expressed Genes (DEGs) were selected on the basis of an increase/decrease of 1.3-fold or greater and a false discovery rate corresponding to a p-value cut-off of 0.05. We identified 530 transcripts that were significantly increased and 403 transcripts that were significantly decreased in arsenite treated *ATF4+/+* neurons compared to *ATF4-/-* neurons (Figure 2C). As expected, amongst the DEGs we identified ATF4 itself as well as several known ATF4 responsive transcripts including DDIT3, SLC7A11, and ASNS. Molecular pathway analysis was then performed on the DEGs using Enrichr analysis package (*68*) which identified 8 pathways that were significantly enriched including the Unfolded Protein Response (UPR) consistent with the known central role of ATF4 in the UPR (*28*). Interestingly, mTOR Signaling and PI3K/AKT/mTOR Signaling pathways were identified amongst the most highly enriched ATF4 responsive pathways (Figures 2D and 2E). This finding is intriguing given the established role of mTOR and PI3K/AKT/mTOR signaling in the regulation of neuronal plasticity, metabolism and survival (*54*).

**Figure 2.**
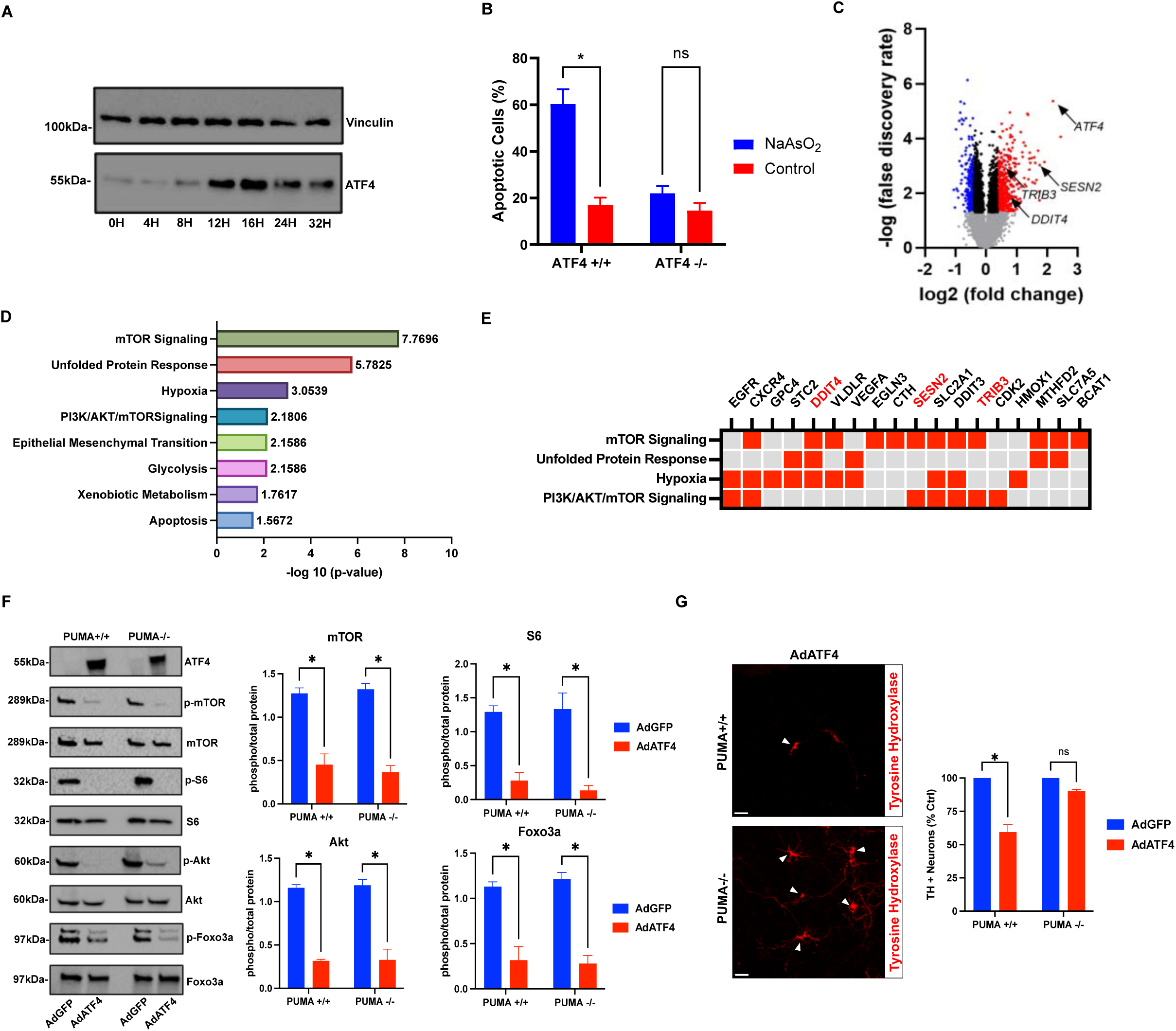
Transcriptome analysis identifies mTOR signaling as a key ATF4 regulated pathway in neurons exposed to oxidative/ proteostatic stress. A. Cortical neurons were treated with sodium arsenite (5μM) and ATF4 expression was assessed by western blot at the indicated times. Vinculin was included as a loading control. **B.** Cortical neurons derived from *ATF4+/+* and *ATF4-/-* mice were treated with sodium arsenite (5μM) and the fraction of apoptotic cells was determined at 24 hours (n=3, 2-way ANOVA, *p<0.05). **C.** Cortical neurons derived from *ATF4+/+* and *ATF4-/-* mice were treated with sodium arsenite (5μM) and mRNA was collected at 12 hours and gene expression was analyzed using the Affymetrix GeneChip platform (3 independent experiments). Volcano plot comparing gene expression profiles in arsenite treated wildtype neurons versus ATF4-deficient neurons. DEGs (>1.3-fold change, p<0.05) are identified in red (increased) and blue (decreased). **D.** Ranking of most enriched terms in Molecular Pathway analysis of DEGs. **E.** Hit map depicting ATF4-dependent DEGs common to the most enriched molecular pathways. Red indicates gene implicated in corresponding pathway and amongst ATF4-dependent DEGs, grey indicates gene not implicated in corresponding pathway. **F.** *PUMA*+/+ and *PUMA*-/- cortical neurons were transduced with Ad-ATF4 or Ad-GFP (50-MOI) and protein was harvested after 24 hours and the levels of p-mTOR (S2448), p-S6 (S235/236), p-AKT (S473) and p-FOXO3a (S253) and corresponding total protein was assessed by western blot and quantified by densitometry. Quantification is reported as the ratio of phosphorylated protein/ total protein (n=3; 2way-ANOVA *p<0.05). **G.** Mesencephalic neuron cultures derived from *PUMA*+/+ and *PUMA*-/- mice transduced with Ad-ATF4 or Ad-GFP (50-MOI) were fixed after 48 hours and immunostained for tyrosine hydroxylase (TH). Representative images showing increased numbers of residual dopaminergic (TH+) neurons (arrow heads) in *PUMA*-/- cultures as compared to *PUMA*+/+ cultures transduced with Ad-ATF4. Quantification of TH+ neurons is reported relative to TH+ counts in Ad- GFP transduced neurons from sister cultures. Data represents the mean ± SEM and statistical differences were determined by 2-way-ANOVA (n=3; *p<0.05).

Next, to determine whether sustained expression of ATF4 can regulate mTOR activity in neurons we transduced primary mouse neuron cultures with a recombinant adenovirus expressing ATF4 (Ad-ATF4/GFP) or a control vector expressing GFP (Ad-GFP). We assessed the effects of sustained ATF4 expression on mTOR activity by examining the phosphorylation state of mTOR and mTORC1/mTORC2 substrates in cortical neurons transduced with Ad-ATF4 versus Ad-GFP. In its activated complexes mTOR is phosphorylated at serine residue S2448 and neurons transduced with Ad-ATF4 exhibit a significant decrease in p-mTOR (S2448) levels as compared to neurons transduced with the control vector Ad-GFP (Figure 2F). Furthermore, neurons ectopically expressing ATF4 exhibit a significant decrease in phosphorylation of ribosomal protein-S6 (S235/236) associated with the mTORC1 pathway and p-AKT (S473) associated with the mTORC2 pathway (Figure 2F). The mTORC2 substrate AKT negatively regulates the transcription factor FOXO3a via phosphorylation preventing its nuclear translocation and transcriptional activity (*cS*). Consistent with his we found that ATF4 expression also caused a significant reduction in FOXO3a (S253) phosphorylation consistent with FOXO3a activation (Figure 2F). Interestingly, we and others have shown that FOXO3a can regulate the transcriptional induction of the pro-apoptotic Bcl-2 family protein PUMA in neurons suggesting a potential link to neuronal death (*70*, *71*). We and others have previously reported that PUMA mediates dopaminergic neuron death in neurotoxin and α- synuclein preformed fibril models of PD (*34*, *72*). Therefore, to determine whether PUMA mediates DA neuronal death induced by ectopic expression of ATF4 we assessed DA neuron survival in *PUMA*+/+ and *PUMA*-/- mesencephalic neuron cultures. DA neurons constitute ∼5% of mesencephalic cultures and were identified by immunostaining for the DA-specific enzyme tyrosine hydroxylase (TH). As shown in figure 2G, enforced expression of ATF4 significantly reduced the number of dopaminergic (TH+) neurons by approximately 40% in *PUMA+/+* cultures, but did not significantly reduce the number of DA neurons in *PUMA-/-*neuron cultures indicating that PUMA is required for ATF4-mediated dopaminergic neuronal death. We also noted that the ATF4-induced downregulation of mTOR activity was similar in *PUMA+/+* and *PUMA-/-* neurons indicating that mTOR suppression was not the result of neuronal death (Figure 2F). Taken together these results demonstrate that sustained expression of ATF4 is sufficient to downregulate mTORC1 and mTORC2 activity and induce PUMA-mediated dopaminergic neuronal death.

### ATF4 suppresses mTOR activity in neurotoxin and α-synucleinopathy models of PD

We previously demonstrated that the DA neuron selective toxins 6-OHDA and MPP+ induce persistent ATF4 expression and ATF4-dependent neuronal death (*34*). Therefore, we next investigated whether mTOR activity is regulated by endogenous ATF4 induction in PD relevant paradigms. To address this, midbrain neurons derived from *ATF4+/+* and *ATF4-/-* littermates were treated with the neurotoxins 6-OHDA or MPP+ and the levels of p-mTOR (S2448) and p-AKT (S473) were assessed by immunofluorescence in dopaminergic (TH+) neurons. As shown in Figure 3A, the levels of p-mTOR and p-AKT were significantly lower in ATF4+/+ DA neurons as compared to ATF4-/- DA neurons following treatment with either 6- OHDA or MPP+ (Figure 3A, Suppl Figure S1A). Although the neurotoxins MPP+ and 6-OHDA are efficiently taken up through the dopamine transporter and preferentially affect DA neurons (*73*) they can also be taken up albeit less efficiently by other neuronal cell types via several other transporters (*74*, *75*). Consistent with this we have previously demonstrated that 6-OHDA and MPP+ induce ATF4 expression and cell death in both dopaminergic and non-dopaminergic neurons although DA neurons are significantly more sensitive (*34*). Therefore, we further investigated the effects of endogenous ATF4 induction on mTORC1 versus mTORC2 activity by western blot in cortical neuron cultures treated with PD relevant neurotoxins. As shown in Figure 3B, 6-OHDA and MPP+ treated cortical neurons exhibited robust ATF4 expression within 8 hours that persisted for at least 24 hours when neuron survival becomes compromised. The increased expression of ATF4 was paralleled by a significant decrease in p-mTOR (S2448) levels beginning at 8 hours that was maintained until at least 24 hours (Figure 3B and Suppl Fig. S1B). Similarly, we found that 6-OHDA and MPP+ induced a robust and sustained decrease in p-S6 (235/236) and p-AKT (S437) levels by 16 hours consistent with downregulation of both mTORC1 and mTORC2 activity (Figure 3B, Suppl. Fig S1B). Furthermore, 6-OHDA and MPP+ treatments decreased the phosphorylation levels of the AKT substrate p-FOXO3a (S253) consistent with increased activity of this pro-death transcription factor. Importantly, the robust downregulation of mTORC1/2 activity induced by 6-OHDA and MPP+ was abolished in ATF4-deficient neuron cultures indicating that ATF4 is required for neurotoxin mediated mTORC1/2 suppression (Figure 3B and Suppl. Fig. S1B).

**Figure 3.**
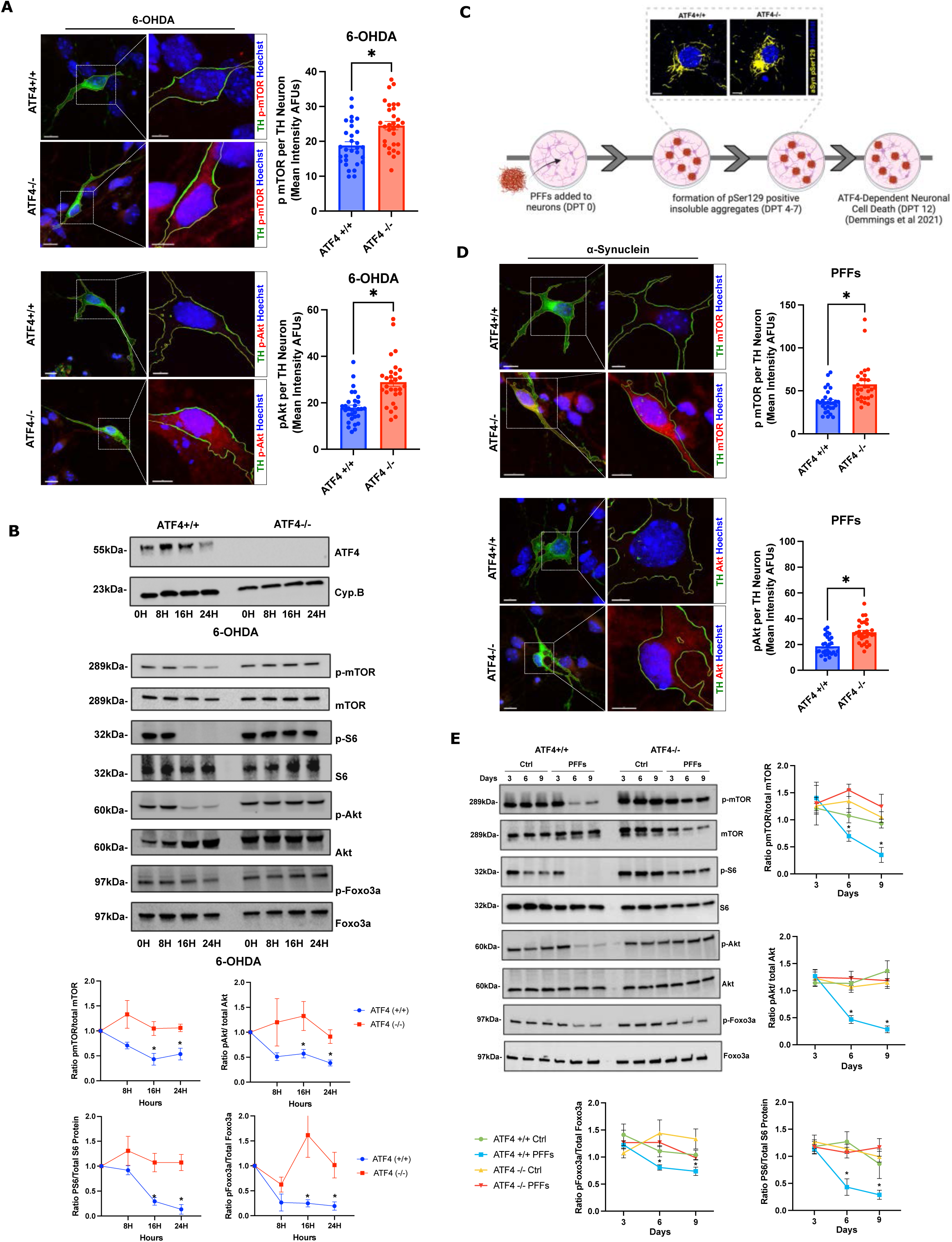
ATF4 regulates neurotoxin and α-synuclein-PFF mediated suppression of mTORC1/mTORC2 activity. A. *ATF4+/+* and *ATF4-/-* mesencephalic neurons treated with 6-OHDA (5μM) were fixed after 24 hours and co-immunostained for tyrosine hydroxylase (TH) and either p-mTOR (S2448) or p-AKT (S473) (scale bar= 15uM). Images on the right have been magnified and the outline of the TH-stained area highlighted to enable visualization of p-mTOR/p-AKT (red) fluorescence (scale bar=5uM). Confocal z-stacked images were acquired and the fluorescence intensity of p-mTOR (S2448) and p-AKT (S473) was quantified in dopaminergic (TH+) neurons from 3 independent experiments (n=29-30 cells per group, t-test, *p<0.05). **B.** *ATF4+/+* an *ATF4-/-* cortical neurons were treated with 6-OHDA (10μM) and protein was harvested at the indicated times and the levels of p-mTOR (S2448), p-S6 (S235/236), p-AKT (S473) and p-FOXO3a (S253) and corresponding total protein was assessed by western blot and quantified by densitometry. Quantification is reported as the ratio of phosphorylated protein/ total protein (n=5; 2way-ANOVA, *p<0.05). **C.** *ATF4+/+* and *ATF4-/-* neurons seeded with α-Syn PFFs (5μg/ml) for 9 days and immunostained for pSer129-α-synuclein. **D.** Mesencephalic neurons derived from *ATF4+/+* and *ATF4-/-* mice were seeded with α-Syn PFFs (5μg/ml) for 9 days and then fixed and co-immunostained for tyrosine hydroxylase (TH) and either p-mTOR (S2448) or p-AKT (S473). Images on the right have been magnified and the outline of the TH-stained area highlighted to enable visualization of p-mTOR/p-AKT (red) fluorescence (left scale bars=15uM, right scale bar=5uM). Confocal z-stacked images were acquired and the fluorescence intensity of p-mTOR (S2448) and p-AKT (S473) was quantified in dopaminergic (TH+) neurons in three independent experiments (n=27-30 cells per group, t-test, *p<0.05). **E.** *ATF4+/+* an *ATF4-/-* cortical neurons were seeded with α-Syn PFFs (5μg/ml) and protein was harvested at the indicated times and the levels of p-mTOR (S2448), p-S6 (S235/236), p-AKT (S473) and p-FOXO3a (S253) and corresponding total protein was assessed by western blot and quantified by densitometry. Quantification is reported as the ratio of phosphorylated protein/ total protein (n=5; 2way-ANOVA, *p<0.05).

We next investigated whether ATF4 regulates mTOR activity in an *in vitro* model of α- synucleinopathy. In this paradigm, neurons take up exogenously added preformed α- synuclein fibrils (α-Syn PFFs) by endocytosis and act as a seed to induce aggregation of endogenous α-synuclein resulting in the formation of intracellular inclusions that cause neuronal dysfunction and cell death (*76*). Synuclein incorporated into pathological inclusions is highly phosphorylated at serine-129 and synuclein inclusions in the PFF model can be readily detected by immunostaining for pSer129-α-synuclein (*77*). We have previously demonstrated that ATF4 expression is upregulated in α-Syn PFF treated DA neurons and cortical neurons and that α-Syn PFF induced neuronal death is attenuated in ATF4-deficient neurons (*34*). As shown in Figure 3C, both *ATF4+/+* and *ATF4-/-* neurons seeded with α-Syn PFFs exhibited pSer129-α-synuclein positive intracellular aggregates detected by immunofluorescence. However, α-Syn PFF induced aggregation caused a significantly greater decrease in p-mTOR and p-AKT levels in *ATF4+/+* DA neurons as compared to *ATF4-/-* DA neurons (Figure 3D). Similarly, α-Syn PFF induced synuclein aggregation in cortical neurons significantly reduced the levels of p-mTOR (S2448), p-S6 (S235/236), p-AKT (S473) and p-FOXO3a (S253) consistent with downregulation of both mTORC1 and mTORC2 activity (Figure 3E). Importantly, α-Syn PFF induced aggregates did not significantly affect mTORC1 or mTORC2 activity in *ATF4-/-* neurons (Figure 3E). Taken together these results demonstrate that sustained induction of ATF4 drives mTORC1 and mTORC2 downregulation in PD neurotoxin and α-synuclein PFF models of PD.

## ATF4 regulates the transcriptional induction of SESN2/DDIT4/Trib3 in neurotoxin and α- synucleinopathy models of PD

To investigate the mechanism by which ATF4 suppresses mTORC1/2 activity we revisited the microarray data (figure 2) to identify transcripts associated with mTOR/AKT signaling pathways that were regulated in an ATF4-dependent manner and that have been reported to inhibit mTOR and/or AKT activity. We identified three transcripts that met these criteria – Sesn2, Trib3, and Ddit4. To confirm that these transcripts are induced in an ATF4-dependent manner we transduced neurons with a recombinant adenovirus expressing ATF4 and assessed Sesn2, Trib3 and Ddit4 mRNA levels by qRT-PCR. Neurons transduced with Ad- ATF4 exhibited a significant increase in Sesn2, Trib3 and Ddit4 mRNA levels compared to neurons transduced with the control vector Ad-GFP suggesting that ATF4 expression is sufficient to induce the expression of these transcripts (Figure 4A). Moreover, neurons treated with MPP+ or 6-OHDA exhibited an ATF4-dependent increase in Sesn2, Trib3 and Ddit4 mRNA levels as determined by qRT-PCR (Figure 4B) and SESN2 and DDIT4 protein levels as determined by western blot (Figure 4C). Of note, we were unable to confirm upregulation of Trib3 protein as we tested several different commercial Trib3 antibodies but none were found to specifically detect Trib3 in mouse neurons. We then examined whether MPP+ and 6-OHDA induce the expression of these transcripts in dopaminergic neurons using multiplex RNAScope *in situ* hybridization combined with TH immunostaining to enable the quantification of these mRNAs in DA neurons. As shown in Figure 4D, the mean number of Sesn2, Trib3 and Ddit4 transcripts detected per DA neuron was significantly higher in MPP+ or 6-OHDA treated *ATF4+/+* DA neurons as compared to *ATF4-/-* DA neurons. Similarly, α-Syn PFF treated wildtype DA neurons also exhibited a significantly greater number of Sesn2, Trib3 and Ddit4 transcripts as compared to α-Syn PFF treated *ATF4-/-* neurons (Figure 4E). Finally, we examined the expression of Sesn and Nipi-3 (orthologues of Sesn2 and Trb3 respectively) in *C. elegans* neurotoxin and α-synuclein overexpression PD models and found that the mRNA levels of both Sesn and Nipi-3 were significantly upregulated in wildtype (N2) but not *Atfs-1-LOF* (Figure 4F-H). These results indicate that ATF4 regulates the transcriptional induction of SESN2, Trib3 and DDIT4 in response to PD neurotoxins and α- Syn PFF induced synuclein aggregation.

**Figure 4.**
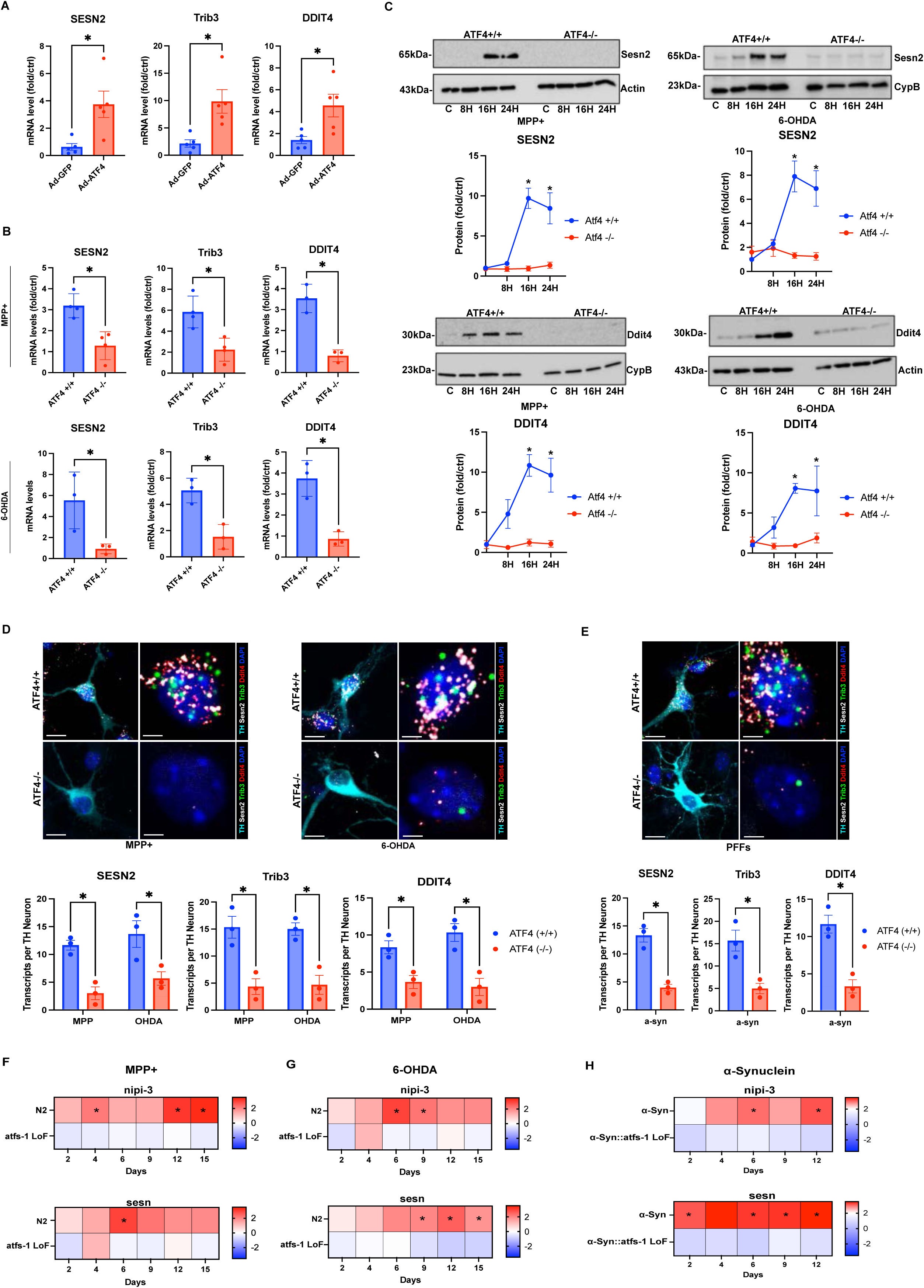
ATF4 regulates the transcriptional induction of SESN2/DDIT4/Trib3 in neurotoxin and α- synucleinopathy models of PD. A. Cortical neurons were transduced with Ad-ATF4 or Ad-GFP (50-MOI) for 24 hours and mRNA levels of Sesn2, Trib3 and Ddit4 were determined by qRT-PCR. Expression is reported as fold increase over untransduced neurons from the same culture (n=5; t-test, *p<0.05). **B.** ATF4+/+ and ATF4-/- cortical neurons were treated with MPP+ (50µM) or 6-OHDA (10µM) for 16 hours and Sesn2, Trib3 and Ddit4 mRNA levels was determined by qRT-PCR. Expression is reported as fold increase over vehicle treated neurons from the same culture (n=3-4; t-test, *p<0.05). **C.** *ATF4+/+* and *ATF4-/-* cortical neurons were treated with MPP+ (50μM) or 6-OHDA (10μM) for the indicated times and SESN2 and DDIT4 protein levels were assessed by western blot and quantified by densitometry and normalized to the loading control Actin or Cyclophilin B (n=3; 2-way-ANOVA *p<0.05). **D.** Mesencephalic neurons derived from *ATF4+/+* and *ATF4-/-* mice were treated with MPP+ (25μM) or 6-OHDA (5μM) for 16 hours and Sesn2, Trib3 and Ddit4 mRNA levels were assessed in dopaminergic neurons using multiplex RNAScope fluorescence *in situ* hybridization combined with tyrosine hydroxylase immunostaining. Representative images showing PD neurotoxin induced expression of Sesn2, Trib3 and Ddit4 mRNA fluorescence puncta in wildtype and ATF4-deficient dopaminergic (TH+) neurons (left scale bar=20uM, right scale bar=5uM). Data represent the mean ± SEM number of transcripts per DA neuron (n=3, 2-way-ANOVA, *p<0.05). **E.** Wildtype and ATF4-deficient mesencephalic neurons were seeded with α-Syn PFFs (5μg/ml) for 9 days and Sesn2, Trib3 and Ddit4 mRNA levels were assessed in dopaminergic neurons using multiplex RNAScope fluorescence *in situ* hybridization combined with tyrosine hydroxylase immunostaining. Data represent the mean ± SEM number of transcripts per DA neuron (n=3, t-test, *p<0.05). **F-H.** RNA was isolated from N2 animals or *atfs-1 LoF* animals treated with MPP+ [100uM] or 6-OHDA [100uM] and *a-syn* or *a- syn::atfs-1LoF* animals at indicated time points. Heat maps represent fold change in gene expression over time as calculated by RT-qPCR. Values were normalized to act-1 housekeeping mRNA and fold change was calculated based on normalized mRNA levels in untreated or wildtype animals at respective time points (n=3, 2way-ANOVA, *p<0.05).

## SESN2/Trib3/DDIT4 co-operate to suppress mTORC1/2 activity and induce PUMA- dependent dopaminergic neuronal loss in neurotoxin and α-synucleinopathy models of PD

SESN2, DDIT4 and Trib3 have been reported to regulate pathways that can suppress mTOR signaling by different mechanisms (*78–81*). Therefore, we investigated whether SESN2, Trib3 and/or DDIT4 induction contribute to mTOR downregulation in neurotoxin and α-Syn PFF models of PD. To address this, we knocked down SESN2, Trib3 or DDIT4 expression using lentiviral-shRNA constructs that co-express target specific shRNAs and GFP under control of the U6 and hPGK promoters respectively (Suppl. Fig. S2A). We first validated the lenti-shRNA constructs in cortical neurons and confirmed high transduction efficiency evident by the high percentage (>90%) of GFP+ neurons (Suppl. Fig. S2B). Furthermore, we confirmed that the 6-OHDA induced expression of SESN2, Trib3 and DDIT4 was efficiently downregulated at the mRNA and protein level in neurons expressing target specific shRNAs but not a non-targeting shRNA (Suppl. Fig. S2C & S2D). To determine whether induction of SESN2, Trib3 and/or DDIT4 is required to suppress mTORC1/2 activity in PD neurotoxin models we transduced cortical neurons with lenti-shRNA vectors specifically targeting these transcripts and then assessed mTORC1/2 activity following treatment with MPP+ or 6- OHDA. As shown in Figure 5A, knockdown of either SESN2, Trib3 or DDIT4 prevented the MPP+ and 6-OHDA induced decrease in p-mTOR (S2448), p-AKT (S437) and p-FOXO3a (S253) levels (Figure 5A). Similarly, knockdown of either SESN2, Trib3 or DDIT4 prevented the α-Syn PFF induced suppression of mTOR (S2448) phosphorylation as well as the phosphorylation of mTORC1 (pS6) and mTORC2 (pAKT, pFoxo3a) substrates (Figure 5A). We then examined the effects of SESN2, Trib3 and DDIT4 knockdown on survival of DA neurons. Knockdown of SESN2, Trib3 or DDIT4 significantly increased DA neuron survival in both neurotoxin (MPP+ and 6-OHDA) and α-Syn PFF treated DA neurons (Figure 5B). Taken together these results indicate that induction of all three of these ATF4 targets (SESN2, DDIT4 and Trib3) are required to efficiently suppress mTORC1/mTORC2 activity and to promote dopaminergic neuronal death.

**Figure 5.**
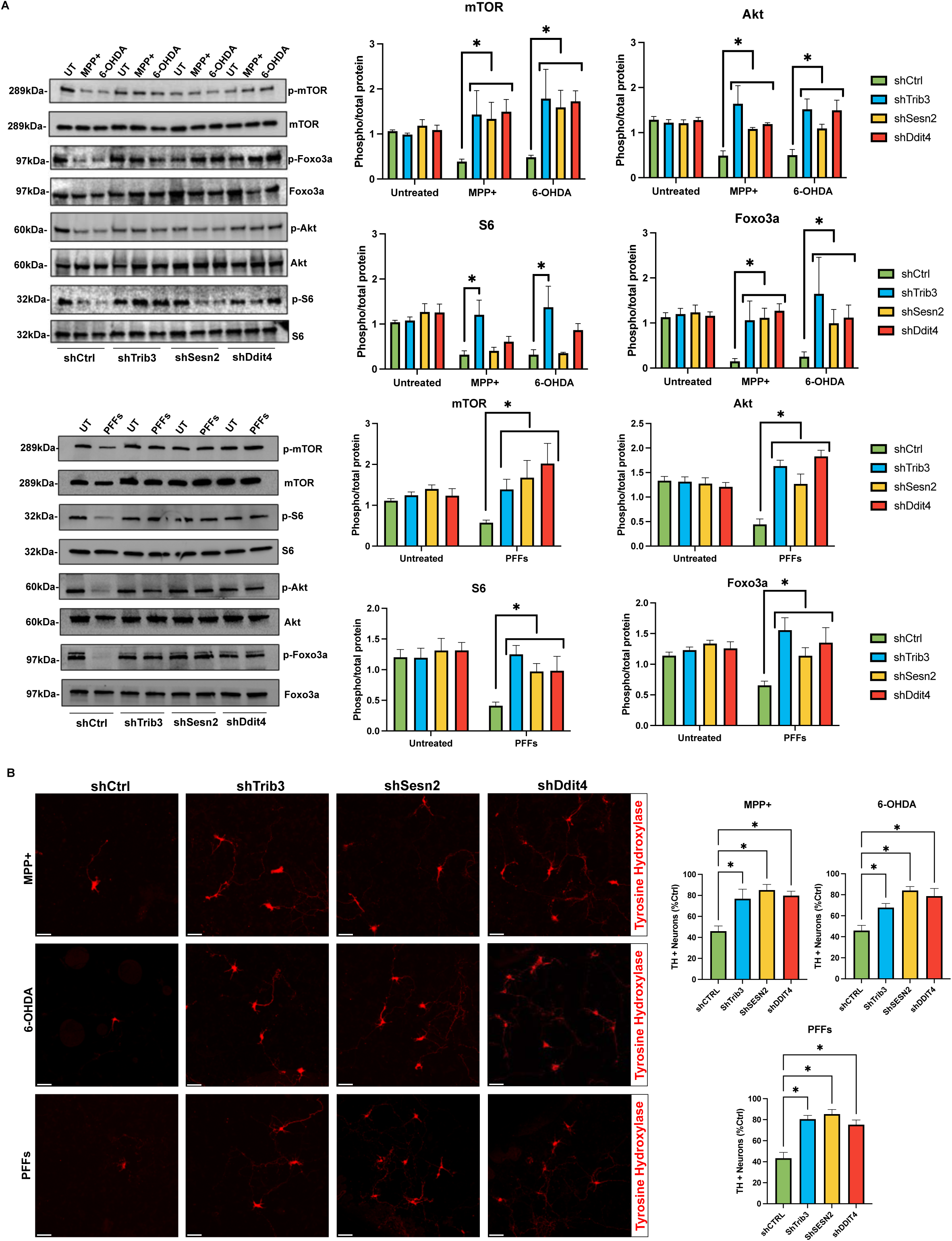
SESN2, DDIT4 or Trib3 knockdown rescues mTOR activity and DA neuron survival in neurotoxin and α-synucleinopathy PD models. A. Cortical neurons were transduced with lenti-shRNA vectors targeting either Sesn2, Trib3, or Ddit4. After 5 days, neurons were treated with MPP+ (50μM) or 6-OHDA (10μM) for 24 hours or seeded with α-Syn PFFs (5μg/ml) for an additional 9 days. The levels of p-mTOR (S2448), p-S6 (S235/236), p-AKT (S473) and p- FOXO3a (S253) and corresponding total protein was assessed by western blot and quantified by densitometry. The ratio of phosphorylated protein/total protein is reported relative to untreated control levels for each shRNA vector (n=5; 2-way-ANOVA, *p<0.05). **B.** Mesencephalic neuron cultures were transduced with lenti-shRNA and after 5 days treated with MPP+ (25μM) or 6-OHDA (5μM) for 48 hours or seeded with α-Syn PFFs (5μg/ml) for an additional 12 days. The number of TH+ neurons was quantified and is reported relative to TH+ counts in vehicle treated neurons from the same culture. Data represents the mean ± SEM and statistical differences were determined by ANOVA (n=4; *p<0.05).

To determine whether expression of SESN2, DDIT4 and Trib3 is sufficient to downregulate mTOR activity we transduced neurons with recombinant viral vectors expressing SESN2, DDIT4 or Trib3 alone or in different combinations. As shown in Figure 6A, the ratio of p-mTOR(S2448)/mTOR was only significantly decreased when all three proteins were co-expressed in neurons (Figure 6A). Moreover, co-expression of SESN2/Trib3/DDIT4 significantly reduced p-S6(S235/236), p-AKT (S473) and p-Foxo3a (S253) levels consistent with suppression of both mTORC1 and mTORC2 activity (Figure 6B). Consistent with the effects on mTORC1/2 activity, we found that DA neuron survival was only significantly decreased when all three proteins (SESN2, DDIT4 and Trib3) were co-expressed in mesencephalic neuron cultures (Figure 6C). Since we and others have previously demonstrated that the pro-apoptotic Bcl-2 family protein PUMA is a key effector of DA neuron death induced by PD neurotoxins and α-Syn PFFs (*34*, *72*, *82*) we sought to determine whether mTORC1/2 suppression is sufficient to induce PUMA expression and DA neuronal loss. To address this, we treated neurons with the selective and potent dual mTORC1/mTORC2 inhibitor OSI-027 (*83*). We confirmed that OSI-027 potently reduced p-mTOR levels in DA neurons (Supplemental Figure S3A) and downregulated mTORC1 and mTORC2 activity in cortical neurons as determined by western blotting of p-mTOR, p-S6 and p-AKT (Supplemental Figure S3B). Therefore, we investigated whether inhibition of mTORC1/mTORC2 activity is sufficient to induce PUMA expression in neurons. As shown in Figures 6D and 6E, OSI-027 significantly increased PUMA expression in DA neurons as determined by immunofluorescence and in cortical neurons as determined by western blot. Furthermore, we found that mTORC1/2 inhibition by OSI-27 was sufficient to decrease DA neuron survival in a dose-dependent manner and that DA neuron loss was significantly attenuated in *PUMA-/-* cultures as compared to *PUMA+/+* cultures (Figure 6F). These results demonstrate that dual inhibition of mTORC1 and mTORC2 is sufficient to induce PUMA expression and PUMA-mediated dopaminergic neuronal death. Finally, we found that co-expression of SESN2/Trib3/DDIT4 caused a significantly greater loss of *PUMA+/+* DA neurons as compared to *PUMA-/-* DA neurons indicating that the pro-apoptotic protein PUMA drives DA neuronal death (Figure 6G). It was also noted that OSI-27 and co-expression of SESN2/DDIT4/Trib3 suppressed mTORC1 and mTORC2 activity to a similar extent in wildtype and PUMA-deficient neurons demonstrating that mTORC1/2 suppression precedes PUMA induction and neuronal death. Taken together these results indicate that the co-induction of SESN2, DDIT4 and Trib3 by ATF4 is necessary and sufficient to suppress mTOR activity and induce dopaminergic cell death in neurotoxin and α-syn PFF models via the induction of the pro-apoptotic BCL-2 family protein PUMA.

**Figure 6.**
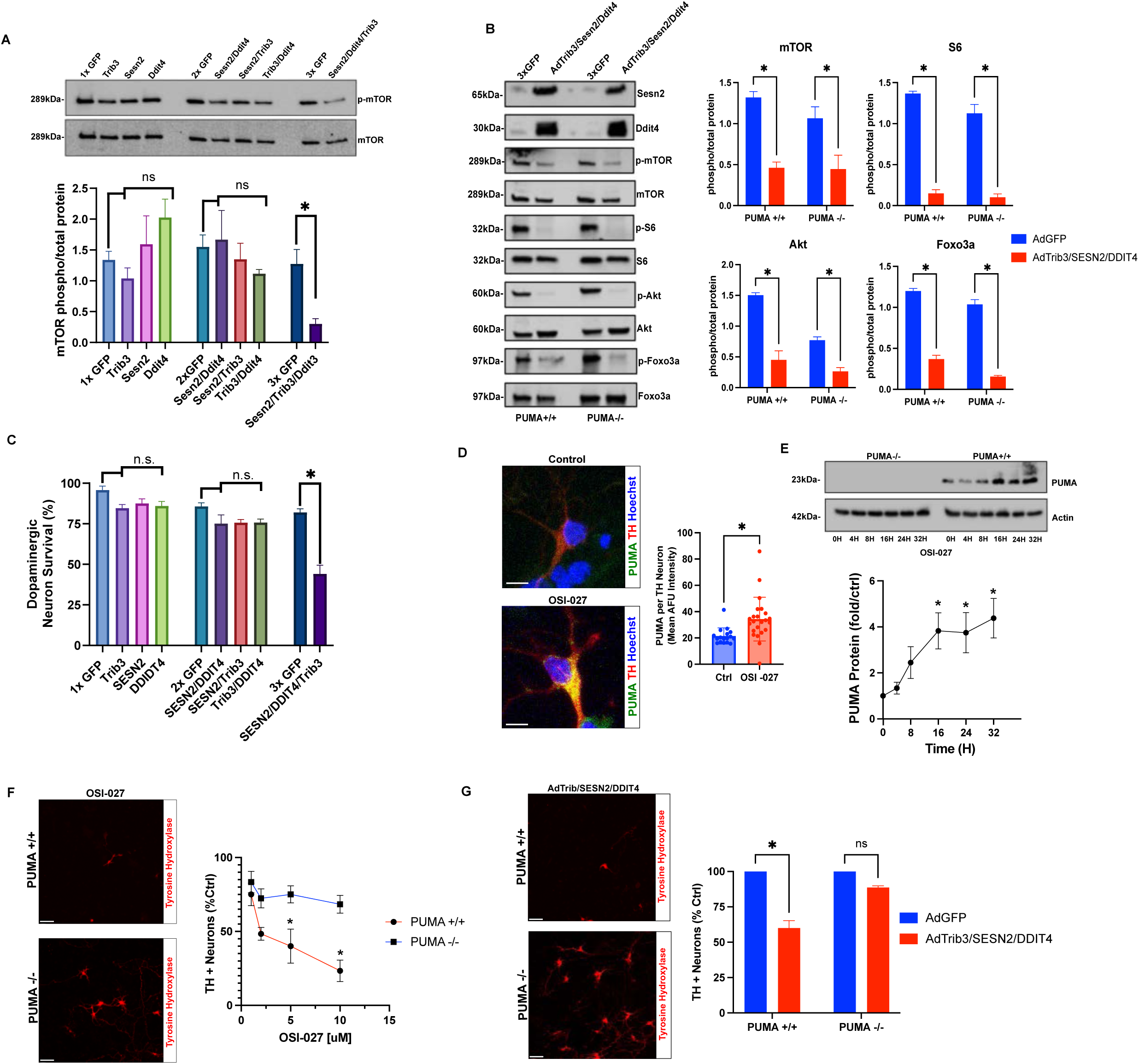
Chronic suppression of mTOR activity by SESN2/Trb3/DDIT4 is sufficient to induce PUMA- mediated dopaminergic neuronal death. A. Cortical or neurons were transduced with recombinant adenoviral vectors (50-MOI) expressing GFP alone or co-expressing GFP and either SESN2, DDIT4 or Trib3 alone or indicated combinations for 24 hours and p-mTOR(S2448)/total mTOR was assessed by western blot and quantified by densitometry (n=3; ANOVA, *p<0.05). **B.** *PUMA+/+* and *PUMA-/-* cortical neurons were co-transduced with Ad-Trib3/Ad- SESN2/Ad-DDIT4 or Ad-EGFP (3X) for 24 hours and p-mTOR (S2448), p-S6 (S235/236), p-AKT (S473) and p-Foxo3a (S253) levels were determined by western blot and quantified by densitometry (n=3; 2-way ANOVA *p<0.05). **C.** Mesencephalic neuron cultures were transduced with indicated vectors and dopamine neuron survival was assessed by TH+ counts at 48 hours. Survival is reported relative to untransduced neurons from the same culture (n=3; ANOVA *p<0.05). **D.** Midbrain neurons were treated with the dual mTOR inhibitor OSI-027 (5μM) for 24 hours and then co-immunostained for tyrosine hydroxylase (TH) and PUMA. Confocal images were acquired and the fluorescence intensity of PUMA was quantified in dopaminergic (TH+) neurons from 3 independent experiments (n=17-23 cells per group; t- test, *p<0.05). **E.** *PUMA+/+* and *PUMA-/-* cortical neurons were treated with OSI-027 (5μM) and PUMA protein levels were assessed by western blot and quantified by densitometry and normalized to the loading control Actin (n=3; ANOVA *p<0.05). **F.** Midbrain neurons derived from *PUMA+/+* and *PUMA-/-* mice were treated with OSI-027 for 48 hours and the number of TH+ cells was counted and reported relative to TH+ counts in vehicle treated neurons from the same culture. Data represents the mean ± SEM and statistical differences were determined by 2-way-ANOVA (n=4; *p<0.05). **G.** *PUMA+/+* and *PUMA-/-* midbrain neuron cultures were transduced with Ad-Trib3/Ad-SESN2/Ad-DDIT4 or Ad-EGFP (3X) and DA neuron survival was assessed by TH+ cell counts after 48 hours (n=3; ANOVA *p<0.05).

## Discussion

The ISR is activated in cells in response to a diverse array of cellular stressors including those relevant to neurodegeneration including oxidative stress, mitochondrial dysfunction and aggregated/unfolded proteins (*24*). While the function of the ISR is to restore homeostasis, chronic activation or hyperactivation of the ISR becomes maladaptive and can cause cell dysfunction and death (*24*). Chronic ISR activation has been implicated in a range of neurodegenerative conditions including Parkinson’s disease (*25*, *27*). Furthermore, genetic deletion and small molecule inhibitors targeting ISR related eIF2a kinases/ eIF2a- phosphorylation have shown therapeutic potential in animal models of FTD, Prion disease, Alzheimer’s disease and Parkinson’s disease (*84*–*90*). However, the role of ATF4, the central transcription factor regulating the ISR response, and the mechanism by which chronic ISR/ATF4 activation becomes maladaptive in neurodegenerative conditions remains unclear. Importantly, in this study we have established that sustained activation of ATF4 promotes dopaminergic neuronal dysfunction and cell death by inducing chronic suppression of mTORC1/2 activity in neurotoxin and α-synuclein models of PD. We previously demonstrated that ATF4 is persistently upregulated in dopaminergic neurons treated with MPP+, 6-OHDA and α-Syn PFFs and promotes DA neuronal loss in these cellular PD models (*34*). In the current study, we incorporated *C. elegans* as a model organism to investigate whether ATFS-1 (ortholog of mammalian ATF4/5) regulates DA neurodegeneration and dopamine function *in vivo*. We demonstrate that similar to mammalian ATF4, ATFS-1 promotes DA neuronal loss in *C. elegans* exposed to PD neurotoxins. Furthermore, we determined that induction of ATFS-1 diminishes the Basal Slowing Rate of animals in neurotoxin and α-synucleinopathy models indicating that the ATF4 ortholog promotes dopamine dysfunction *in vivo* and results in shortened animal life span in this pathological context. Taken together these results suggest that sustained activation of ATF4 in DA neurons under chronic oxidative and proteostatic stress promotes dopaminergic neuronal dysfunction and degeneration. Consistent with our findings, the Caldwell lab has demonstrated that chronic overexpression of ATFS-1 in *C. elegans* is sufficient to induce DA neurodegeneration and decreased dopamine function as assessed by BSR assay (*63*). On the other hand, *C. elegans* strains expressing *pdr-1 LoF* (mammalian ortholog of Parkin) or *Pink-1 LoF* alleles do not exhibit DA neurodegeneration but do exhibit modest DA neuron loss on an *atfs-1 LoF* background suggesting that mild or transient activation of UPR^MT^ ATFS-1 exerts neuroprotective properties (*91*). Overall, these data support a model in which under mild UPR/ISR conditions ATFS-1/ATF4 can effectively restore cell homeostasis and protect neurons, whereas under chronic or hyperactivation conditions ATFS-1/ATF4 activation becomes maladaptive and promotes neuronal dysfunction and degeneration. In future studies it will be important to investigate whether ATF4 affects neurodegeneration in mouse models of PD that have a more complex and disease relevant nigrostriatal network within a glial environment, and would enable the assessment of dopaminergic function in motor behavior paradigms that better represent those affected in human PD.

In this study we establish that ATF4-mediated downregulation of mTOR activity drives DA neuron death in both PD neurotoxin and α-synucleinopathy models. Since mTOR signaling promotes general protein synthesis and metabolic activity, its transient downregulation could be beneficial by permitting selective ISR-mediated adaptive gene expression and decreasing cellular energy demands to facilitate cell recovery. However, sustained deactivation of mTOR would be detrimental as mTOR signaling has been implicated in the regulation of synaptic plasticity, dopaminergic neural transmission and dopamine-dependent behaviours (*42*, *S2*–*S4*) that would be compromised during persistent ISR/ATF4 activation. Consistent with this, numerous studies using genetic and pharmacological approaches have demonstrated the involvements of ISR related eIF2α/eIF2α kinases in synaptic plasticity, long-term memory formation and dopamine-dependent behaviours (*95*– *100*). Furthermore, since mTORC2 regulates the activity of AKT, sustained mTOR deactivation would result in the loss of AKT pro-survival signaling. Consistent with this we demonstrate that p-AKT (S473) levels are markedly reduced in parallel with mTOR in both neurotoxin and α-Syn PFF paradigms. AKT negatively regulates the transcription factor FOXO3a via phosphorylation preventing its nuclear translocation and transcriptional activity (*cS*) and we demonstrate that FOXO3a (S253) phosphorylation is also markedly reduced in an ATF4- dependent manner in neurotoxin and α-Syn PFF models. It has previously been reported that FOXO3a can promote the transcriptional induction of the pro-apoptotic Bcl-2 family protein PUMA (*70*, *71*) and we and others have shown that PUMA-deficient dopaminergic neurons are resistant to PD neurotoxin and α-Syn PFF induced cell death (*34*, *72*, *82*). Furthermore, we show that ectopic expression of ATF4 or pharmacological inhibition (OSI-027) of mTOR results in AKT deactivation, FOXO3a dephosphorylation (activation), PUMA induction and PUMA-dependent DA neuron loss suggesting a potential mechanistic link to neuronal death. In line with our findings, it was shown that transduction of dopamine neurons with AAV expressing a constitutively active form of Rheb, a key activator of mTOR signaling, attenuates nigrostriatal degeneration in 6-OHDA lesioned mice (*101*). PTEN is a negative regulator of AKT and mTOR activity, and it has been shown that specific deletion of Pten in dopaminergic neurons protected DA neurons and enhanced DA-dependent behavioural functions in 6-OHDA lesioned mice (*102*, *103*). However, other studies have reported beneficial effects of specific inhibition of mTORC1 with Rapamycin or its analogs in rodent models of synucleinopathy (*54*). In these studies, Rapalogs are proposed to function by mitigating mTORC1 mediated inhibition of the autophagy activator ULK1 to enhance the autophagic clearance of α-synuclein aggregates. However, in a recent study, it was reported that selective inhibition of mTORC1 protects dopaminergic neurons in synucleinopathy models independent of its effects on autophagy (*104*). In our studies we found that ATF4 suppressed both mTORC1 and mTORC2 activity, but it is unclear whether the death promoting activity of sustained ATF4 activation is mediated by mTORC1, mTORC2 or requires suppression of both complexes. Future studies to selectively inhibit or restore activity of mTORC1 versus mTORC2 to determine complex specific contributions to neurodegeneration would be highly valuable from a therapeutic perspective.

To investigate the mechanism by which ATF4 mediates mTOR downregulation we screened our microarray for DEGs that have previously been reported to repress mTOR/AKT activity and identified DDIT4 (aka REDD1/RTP801), SESN2 and Trib3 (*80*, *81*, *105*, *106*). Interestingly, it has been reported that TRIB3 and DDIT4 (RTP801) expression is elevated in post-mortem brain tissue of PD patients and in cellular models of PD (*107*, *108*). We demonstrate that Trib3, SESN2, and DDIT4 are transcriptionally induced in an ATF4-dependent manner in PD neurotoxin and α-synucleinopathy paradigms. Furthermore, we demonstrate that knockdown of either Trib3, SESN2 or DDIT4 expression prevents neurotoxin and α-Syn PFF induced mTOR deactivation and DA neuron loss. This suggests that these three proteins co-operate to downregulate mTORC1and mTORC2 activity and consistent with this we found that adenoviral mediated co-expression of all three proteins (Trb3/DDIT4/SESN2) is required to significantly reduce mTOR activity and DA neuron survival. SESN2 and DDIT4 have been shown to inhibit mTORC1 by distinct mechanisms. SESN2 has been shown to inhibit mTORC1 by inhibiting GATOR2 and preventing Rag-mediated activation of mTORC1 (*106*). DDIT4, on the other hand, has been reported to inhibit mTORC1 by binding 14-3-3 proteins preventing the dissociation of the TSC1/TSC2 complex and preventing Rheb mediated activation of mTORC1 (*109*). Trib3 has been shown to bind and inhibit both AKT and mTORC2 (*80*, *81*). However, while mTORC1 and mTORC2 can be differentially regulated, crosstalk between these pathways has been well documented and for example the mTORC2 substrate AKT is known to inhibit the TSC1/TSC2 complex to promote mTORC1 activity (*110*). Therefore, we propose a model in which SESN2, DDIT4 and Trib3 inhibit mTOR through distinct pathways and thereby prevent crosstalk and compensatory mechanisms to persistently suppress both mTORC1 and mTORC2 signaling.

In summary, we have established a critical molecular pathway that regulates DA neuron survival in PD neurotoxin and α-synucleinopathy models providing novel therapeutic targets for the treatment of PD. Specifically, we demonstrate that chronic ISR/ATF4 activation induces Trib3/DDIT4/SESN2 expression that co-operate to suppress mTOR signaling resulting in PUMA-dependent DA neuronal death. Furthermore, we suggest that this pathway may have broader implications as chronic ISR activation and mTOR dysregulation are common features of several neurodegenerative diseases.

## Materials G Methods

### Animals

All animal procedures were performed as per guidelines set by the Animal Care Committee at Western University, in accordance with the Canadian Council on Animal Care. ATF4-null mice were provided by Dr. Tim Townes (University of Alabama) and PUMA-deficient animals were a gift from Dr. Andreas Strasser (WEHI, Melbourne, Australia). Animals were maintained on a C57/BL6 background, and heterozygous mice were crossed to obtain wildtype and knockout littermates for experimentation. Animals were genotyped as previously described (*111*, *112*).

### Primary Cortical and Mesencephalic Neuron Cultures

To generate primary neuron cultures pregnant dams were sacrificed and cortical neurons were dissociated from embryonic day 14.5-15.5 males and females. Following tissue processing, neurons were cultured in Neurobasal/B27 plus system (ThermoFisher, #A35829- 01) supplemented with Glutamax (ThermoFisher, #35050-061) and Antibiotic/Antimycotic (ThermoFisher, #15240062) as previously described (*113*). To generate dopaminergic neuron cultures, the ventral mesenchephalon was dissected from E14.5 -15.5 embryonic mice and neurons were cultured in the media described above in a protocol modified from Gaven and colleagues (*114*).

### Drug Treatments

Drug treatments were initiated in cortical and mesencephalic neuronal cultures at 7 DIV unless specified otherwise. Stock solutions of 1-methyl-4-phenylpyridiniumiodide (MPP+; Millipore Sigma #D048), 6-hydroxydopamine hydrobromide (6-OHDA; MilliporeSigma #H116), and Sodium Arsenite (NaAsO2; Millipore Sigma #S7400) were prepared in ddH2O. Stock solutions of OSI-027 (SelleckChem #S2624) were prepared in DMSO. Drugs were diluted in neuron culture media immediately before administration to cultures at indicated concentrations.

### Generation of human alpha-synuclein preformed fibrils

Human synuclein preformed fibrils were generated from recombinant human alpha synuclein monomers (Proteos #RP-003) as previously described (*34*), based on the protocol of Volpicelli-Daly and Colleagues (*115*). In brief, monomers were resuspended in modified PBS and were shaken at 1000rpm at 37°C for 7 days. Beta-sheet confirmation was determined for each batch of PFFs by Thioflavin-T staining (Sigma #T3516). Prior to adding to neurons, PFFs were sonicated using a Fisher Scientific Sonicator Dismembrator (Model 100) at 10% power using 0.5 second pulsation for a minimum of 30 seconds. Sonicated PFFs were then added to neuronal cultures at DIV 7 at a concentration of 5ug/mL for the indicated amount of time.

### Quantitative RT-PCR

RNA was isolated via Trizol extraction as per the manufacturer’s instructions (Invitrogen # 15596018) and 40 ng was loaded into QuantiNova reverse transcription (RT)-PCR mix along with target specific primers (Supplemental Table 1) as per provided protocol (Qiagen #208352). qRT-PCR was performed on a CFX Connect Real Time System (Biorad) and changes in gene expression were determined by the Δ(ΔCt) method using S12 transcript for normalization. Values for qRT-PCR are reported as fold increase in mRNA levels in treated samples relative to untreated controls for transcript of interest.

### Western Blot Analysis

To obtain whole-cell lysates, neurons were incubated in RIPA lysis buffer (Millipore Sigma, #R0278) supplemented with PhoSTOP phosphatase inhibitor cocktail (Roche # 4906845001) and cOmplete protease inhibitor cocktail (Roche # 04693132001) on ice for 30 minutes and the soluble fraction was collected via centrifugation. Protein concentration was determined by BCA assay (ThermoScientific, #23225) and 40ug of protein was separated by SDS-PAGE and transferred onto PVDF membranes. Membranes were incubated in primary antibody (Supplemental Table 2) overnight at 4°C. Membranes were then washed in TBST before the addition of the secondary antibody conjugated to HRP at room temperature for 2 hours. All antibodies were prepared in 5% BSA in TBST (10 mM Tris, 150 mM NaCl, 0.1%Tween 20). Membranes were then washed and developed via enhanced chemiluminescence (BioRad #1705061). Chemiluminescence signal was detected with a Bio-Rad ChemiDoc MP Imaging System and densitometric measurements were determined using ImageLab software (Bio-Rad). Protein levels were normalized to loading control and reported as the ratio of phosphorylated to total protein.

### Apoptotic cell counts

Neuronal apoptosis was calculated by visualizing nuclear morphology in Hoechst 33342 stained cells as described previously (*34*). Briefly, neurons were fixed in Lana’s Fixative (4% paraformaldehyde, 0.2% picric acid) for 30 minutes and stained with Hoechst 33342 (Invitrogen #H1399) at a concentration of 0.5 µg/mL for 10 minutes. Apoptotic nuclear morphology was quantified by an individual blinded to the treatments and is reported as the fraction of cells exhibiting pyknotic and/or fragmented nuclei containing condensed chromatin in a minimum of five randomly selected fields (minimum of 500 cells per treatment).

### Immunoffuorescence and Quantification

Neurons were washed with ice cold PBS containing Na_3_VO_4_ (1 mM; Millipore Sigma #S-6508)) and NaF (25mM; Millipore Sigma # S-6776) and then fixed in Lana’s fixative (4% paraformaldehyde, 0.2% picric acid) for 30 minutes. Neurons were then permeabilized with 0.3% Triton X-100 and blocked in 4% normal goat serum for 1 hour and incubated with primary antibodies (Supplemental Table 2) in 0.3% Triton-X100, 2% normal goat serum in PBS overnight at 4 °C. The following day cells were washed with PBST and incubated with appropriate Alexa Fluor (Invitrogen) conjugated secondary antibodies in 2% goat serum/0.3% Triton-X100 in PBS for 2 hours at room temperature. Neurons were then washed in PBS-T and nuclei stained with Hoechst prior to mounting on SuperFrost glass coverslips (FisherBrand #22-037-246) using Prolong Gold mounting media (ThermoFisher #P36982). Images were acquired on a Leica SP8 Confocal microscope and TH+ neurons were randomly identified for imaging using LAS X Navigator software (Leica Microsystems). Confocal z-stacked images of dopaminergic neurons (TH+) were captured at 40X magnification using optimized and consistent laser power and constant gain with excitation/emission profiles appropriate for the secondary antibody of interest. Imaris software (Oxford instruments) was used to quantify fluorescence signal intensity, specifically in dopaminergic neurons. Briefly, confocal images were converted to Imaris (.ims) files and surfaces around dopaminergic neurons were created utilizing the Imaris surface creation tool based on the absolute intensity of TH signal. Given the expected activities of the proteins of interest in either the cell body or nucleus and to not bias the reading of mean intensity by the number of projections included in the surface, Regions of interest (ROI) were set around the cell body marked by the brightest intensity of TH signal and the surface utilized for analysis of mean fluorescence intensity was generated within this ROI. Mean fluorescence intensity was then measured as the average pixel intensity of the signal of interest within the cell body and is reported as arbitrary fluorescence units (AFU). A minimum of 10 dopaminergic neurons in a minimum of three independent experimental replicates (>30 cells per treatment group) was assessed for each treatment condition.

### Tyrosine Hydroxylase Cell Quantification

Mesencephalic neuron cultures plated on coverslips in 4-well plates were immunostained for tyrosine hydroxylase as described above. Neurons were visualized using a Leica SP8 confocal microscope and all TH+ neurons on coverslips were identified using the LAS X Navigator scanning software package (Leica microsystems). The total number of dopaminergic (TH+) cell bodies per coverslip was then scored by an individual blinded to the treatment conditions and survival was determined as the fraction of TH+ cells remaining in treated versus vehicle treated cultures and reported as a percentage.

### Dual TH immunoffuorescence and Multiplex RNAScope ffuorescence in situ hybridization

Mesencephalic neurons cultured on coverslips were fixed for 20 min in Lana’s fixative (4% paraformaldehyde, 0.2% picric acid) and dehydrated in ethanol and then coverslips were removed from culture dishes and adhered to microscope slides (FisherBrand #1255015) with xylene based adhesive and a surrounding hydrophobic barrier was created. Cells were then rehydrated and RNAScope multiplex fluorescence *in situ* hydridization was completed as per manufactures instructions (Advanced Cell Diagnostics ACD #323136). Briefly, slides were incubated in hydrogen peroxide for 10 min then washed with PBS. Cells were then permeabilized with RNAScope Protease III for 10 min at room temperature and washed in PBS. Probe hybridization for Trib3 (ACD #506301), SESN2 (ACD # 575751), and DDIT4 (ADC #468551) was carried out for 2 h at 40 °C in a HybEZ II humidity-controlled oven (ACD #321720). Probes assigned to specific channels underwent sequential amplification and were then recognized via incubation with TSA vivid fluorophores (Tocris). After the completion of RNAscope assay, slides were then stained for tyrosine hydroxylase using the standard protocol described above. Slides were then incubated with DAPI for 30 seconds and mounted on glass slides with Prolonged Gold media (ThermoFisher #P36982). Cells were visualized using a Leica SP8 confocal microscope and TH+ cells were randomly identified using LAS X Navigator Software and then z-stacked images were automatically acquired from each TH+ neuron at optimized excitation/emission windows for each target of interest. Images for each transcript were individually overlayed on TH/DAPI images and the number of puncta in the soma/nucleus of each TH+ cell was counted manually by an individual blinded to the treatment. A minimum of 10 dopaminergic neurons in each treatment group was scored by an individual blinded to the treatment in at least three independent experiments (>30 cells per treatment group).

### Adenoviral Construct Generation

Adenoviral vectors expressing GFP or GFP-P2A-ATF4 were generated as previously described 33. Shuttle plasmids for the generation of recombinant adenoviral vectors expressing Trib3, DDIT4 and SESN2 were constructed from pcDNA-Trb3 (Addgene #131157), pCMS-eGFP- RTP801 (Addgene #65057) and pGEX-Sesn2 (Addgene #61873) respectively in a two-stage cloning process. In the first stage Trib3, DDIT4 and SESN2 cDNAs flanked by 5’-BamHI and 3’- EcoRI restriction sites were generated by PCR amplification (Trib3, DDIT4) or restriction enzyme digest (SESN2) of the base plasmids and inserted into the BamHI/EcoRI MCS of the pUltra-EGFP plasmid (Addgene #24129) to enable bicistronic expression of EGFP. Coning primer sequences are available upon request. In the second step, the EGFP-P2A-Trib3, EGFP-P2A-DDIT4, and EGFP-P2A-SESN2 cassettes were inserted into the MCS of the pShuttle-CBA-SV40 vector to enable production of recombinant adenovirus in HEK 293A cells using the protocol described by Luo and colleagues with modifications (*116*). Briefly, pShuttle constructs were linearized with PmeI and electroporated into BJ5183-AdEasier-1 cells (Addgene#16399) containing the pAdEasy-1 adenovirus backbone. We next identified positive recombination by PacI digest and positive clones were amplified in Stbl2 bacterial cells (ThermoFisher #10268019). Recombinant AdEasy-shuttle plasmids were digested with PacI and transfected into HEK293A cells (ATCC CRL-1573) using Lipofectamine (ThermoFisher #11668027) to generate adenovirus particles. Following this, cells were harvested after 14 days and crude viral lysate was obtained and amplified in successive rounds in HEK293A cells. Amplified viral lysates were then purified by CsCL gradient centrifugation and desalting by dialysis using Slide-A-Lyzer cassettes with 10 000 MW cut off (Thermofisher #66380). Finally viral particles were filtered through 0.22uM PES syringe filters. Adenovirus preparations were then tittered in HEK293 cells and MOI was calculated based on GFP expression.

### Lentivirus expressing shRNA

Lentiviruses expressing shRNAs specifically targeting Trib3, SESN2 and DDIT4 were purchased from Vectorbuilder (shCTRL VB# VB010000-009mxc, shTrib3 VB# VB230127- 1198jed, shSESN2 VB# VB230127-1195msp, shDDIT4 VB# VB230127-1197gke). The lentiviral vectors were designed to express specific shRNA and EGFP under control of the U6 promoter and hPGK promoter, respectively. Primary neurons were transduced with 10 MOI of indicated lentivirus on DIV 3 and monitored for GFP expression to determine transduction efficiency, by day 5-post-transduction over 90% of neurons were GFP positive and experiments were initiated.

### C. elegans strains and maintenance

*C. elegans* were maintained as previously described (*117*) on nematode growth medium (NGM) agar plates at 20 C in a ReptiPro 6000 incubator unless otherwise stated. Animals were fed with a bacterial lawn of live OP50 *E. coli* obtained from Caenorhabditis Genetic Center (CGC), University of Minnesota. The following strains were provided by the by the CGC: wildtype (N2), *unc-54p::α-synuclein::YFP + unc-119(+)* (NL5901), *dat-1p::GFP* (BZ555), *atfs-1 LoF* (VC3201). Prior to all experimental procedures worms were synchronized via alkaline hypochlorite bleach solution (0.8% NaOCl, 0.5 M NaOH), in which only embryos are persevered. Embryos were then transferred to and maintained in S-complete media (S-basal (NaCl, 1 M potassium phosphate pH 6.0), 1 M potassium citrate (tripotassium citrate, citric acid monohydrate, 5M KOH), trace metal solution (Na2EDTA, FeSO4, 1M CaCl2, 1M MgSO4•7H2O, MnCl2•4H2O, ZnSO4•7H2O, CuSo4), 1 M CaCl2, 1 M MgSO4), OP50, streptomycin (Fisher #11860038), and 0.6 mM 5-Fluoro-2’-deoxyuridine (FUDR; Sigma #F0503) to prevent generation of new progeny.

### Generation of C. elegans Crosses

Using transgenic strains sourced from CGC, we generated two specific crosses 1) *atfs-1 LoF::dat-1:GFP* and 2) *atfs-1 LoF::unc-54p::a-syn::YFP*. Briefly animals from each genotype were crossed to generate an F1 offspring. Animals from the F1 cross were plated as single worms in individual wells to facilitate population expansion via hermaphrodite-proliferation for 1-2 weeks. The F1 progeny was then screened for either *dat-1:GFP* or *unc-54p::a- syn::YFP* based on the presence of the fluorescence reporter. Animals exhibiting the appropriate fluorescence reporter were then genotyped and animals heterozygous for the *atfs-1 LoF* allele were confirmed via PCR with sequence specific primers for the 881 BP deletion. Next, heterozygous *atfs-1 LoF* animals with the appropriate fluorescence reporter were then crossed to generate the F2 progeny. F2 animals were plated and screened based on fluorescence reporter and confirmed to be homozygous for *unc-54p::a-syn::YFP*. F2 animals were then subsequently confirmed to be *atfs-1 LoF* homozygous animals via genotyping. Desired genotypes were then maintained as described above.

### C. elegans drug treatment

The neurotoxins 1-methyl-4-phenylpyridinium iodide (MPP+; Millipore Sigma #D048) or 6- hydroxydopamine hydrobromide (6-OHDA; Millipore Sigma #H116) were diluted directly into a liquid culture consisting of Adult L4 worms to achieve the indicated final concentration.

### C. elegans Lifespan Assay

Lifespan assays were conducted in liquid S-complete cultures in 96-well plates as described (*118*). Animals were visualized using an inverted-light microscope and scored as live based on movement and light response reflex. Immediately before treatment, a baseline count of live *C. elegans* was recorded, and *C. elegans* were scored at indicated time intervals for approximately 30 days or until all worms had died. A minimum of 120 worms over 4 independent experiments was analyzed for each treatment group.

### Isolation of C. elegans RNA and Protein

Prior to RNA isolation or protein isolation, *C. elegans* were pelleted and washed several times in M9 buffer (Na_2_HPO_4_, KH_2_PO_4_, NaCl, MgSO_4_·7H_2_O) to limit any bacterial carryover from growth media. RNA was isolated from pelleted animals using Trizol reagent (Invitrogen

\# 15596018) as per manufacturers protocol. qRT-PCR was performed as described above using gene specific primers and act-1 for normalization (primer sequences listed in Supplemental Table 1). To isolate protein, pelleted worms were incubated in RIPA buffer supplemented with 0.1% SDS and 10 uM DTT, phosphatase inhibitor cocktail (Millipore Sigma, #P5726) and protease inhibitor cocktail (Millipore Sigma, #P8340). To ensure efficient lysis, samples were further sonicated with a Sonic Dismembrinator Model 100 (Fischer Scientific) for 4 rounds of six successive 1 second on/ 1 second off pulses at 50% power. Protein quantification, and western blotting with *C. elegans* protein was carried out as described above.

### Scoring dopaminergic neuron survival in C. elegans

Transgenic *C. elegans* (*dat-1:GFP*) expressing GFP specifically in dopaminergic neurons were collected from experimental groups and centrifuged to pellet worms. Pellets were then washed to minimize bacterial carryover and then worms were anesthetized and mounted on coverslips based on a protocol adapted from Kelley et al., 2017 (*119*). Briefly, worms were anesthetized via 0.2% (wt/vol) levamisole and 2% (wt/vol) tricaine diluted in M9 buffer for 30 minutes. Following anesthetization, worms were plated on 5% agar pads on microscope slides and visualized on a fluorescence microscope (IX70 Olympus). To quantify dopaminergic neuron survival, a minimum of 10 animals from each treatment group were randomly selected and the number of intact dopaminergic neurons (GFP+ cell bodies) was counted in each animal by an evaluator blinded to the treatment conditions. Dopaminergic neuron survival was determined by the fraction of counted GFP+ soma (dopaminergic neurons) out of the 6 CEP/ADE dopaminergic neurons for each animal and reported as a percentage.

### C. elegans Basal Slowing Response

Worms from indicated experimental groups were split into two groups and transferred onto 60mm NGM agar plates with or without an OP50 bacterial lawn food source according to a protocol modified from Sawin et al., 2000 (*64*). *C. elegans* locomotion was then recorded by a Stingray F-504 B CCD monochrome camera (Allied Vision Tech) equipped with a Navitar Zoom 7000 lens, and darkfield illuminator (Schott) at 9 frames per second for 60 seconds/540 frames using MATLAB image acquisition tool. Recordings were then analyzed and the speed of individual worms were quantified using the ImageJ wrMTrck plugin (available at: http://www.phage.dk/plugins/wrmtrck.html) created by Jesper Pedersen. The average speed of each experimental group was then used to calculate basal slowing rate (BSR) using the following equation: BSR= (speed in the presence of food/speed in the absence of food) x 100. Approximately 50 worms were scored in each experimental group.

### Affymetrix Microarray Transcriptome Analysis

Total RNA was isolated from vehicle or NaAsO_2_ treated *ATF4+/+* and *ATF4-/-* cortical neuron cultures (3 independent experiments) by Trizol extraction as per manufacturer’s instructions (Invitrogen #15596018). RNA quality (RIN scores >9) was confirmed using the Agilent 2100 Bioanalyzer (Agilent Technologies Inc., Palo Alto, CA) and the RNA 6000 Nano kit (Caliper Life Sciences, Mountain View, CA). All sample labeling and GeneChip processing was performed at the London Regional Genomics Centre (Robarts Research Institute, London, Ontario, Canada; http://www.lrgc.ca). Briefly, single stranded complimentary DNA (sscDNA) was prepared from 100 ng of total RNA as per the Affymetrix GeneChip WT PLUS Reagent Kit (Affymetrix, Santa Clara, CA). Total RNA was first converted to cDNA, followed by *in vitro* transcription to make cRNA. 5 ug of single stranded cDNA was synthesized, end labeled and hybridized, for 16 hours at 45°C, to Mouse Gene 2.0 ST arrays. All liquid handling steps were performed by a GeneChip Fluidics Station 450 and GeneChips were scanned with the GeneChip Scanner 3000 7G (Affymetrix, Santa Clara, CA) and probe level data was generated using Affymetrix Command Console v3.2.4. Probes were summarized to gene level data in Partek Genomics Suite v6.6 (Partek, St. Louis, MO) using the RMA algorithm (*120*). Partek was used to determine gene level ANOVA p-values and fold changes and Differentially Expressed Gene (DEG) lists were created using a filter of 1.3-fold change and p-value of < 0.05. Further pathway and gene-enrichment analysis of DEGs was performed using the EnrichR analysis package (*68*). We the used the Enrichment Analysis visualizer appyter linked to the EnrichR gene set to perform molecular pathway analysis of the gene list (all genes greater than 1.3-fold change and p-value<0.05) based on the MSigDB Hallmark 2020 gene set library. Full gene expression data sets available at https://doi.org/10.5061/dryad.xgxd254r2.

### Statistical analysis

Data are reported as mean ± standard error of the mean (SEM) from at least three independent experiments. Sample size was estimated using pilot data and power analysis method. Statistical analyses were performed using Graph-Pad Prism version 8.2 for Mac (GraphPad Software, USA). Statistical significance was determined using two-tailed paired or unpaired Student’s t-test for individual comparisons, one-way ANOVA followed by Tukey post-hoc test and two-way ANOVA followed by Sidak post-hoc tests for multiple comparisons and differences were considered significant at P < 0.05. In Figure Legends, ‘n’ value indicates the number of independent experiments and/or the number of mice of each genotype from which cultures were prepared in independent experiments.

## Resource Availability

*Lead Contact:*

Sean Cregan

## Materials availability

All unique reagents generated in this study are available from the lead contact without restriction.

## Data and Code availability

All data reported in this paper will be shared by the lead contact upon request. This paper does not report original code.

Any additional information required to reanalyze the data reported in this paper is available from the lead contact upon request

## Acknowledgements

Canadian Institutes of Health Research grant PJT-178371 (SPC) Parkinson’s Foundation Summer Fellowship (KH)

## Author Contributions

Conceptualization: SPC, MDD Methodology: MDD, EAK, ECT Investigation: MDD (lead), EAK, KH, VC Formal Analysis: MDD, JZ, JK, NC

Visualization: SPC, MDD, ECT Supervision: SPC, SHP, MDD Writing: SPC, MDD **Declaration of Interest**

Authors declare that they have no competing interests

## Supplemental Information

Document S1: Figures S1-S3, Table S1-S2.

---

**Supplemental Figure S1.**
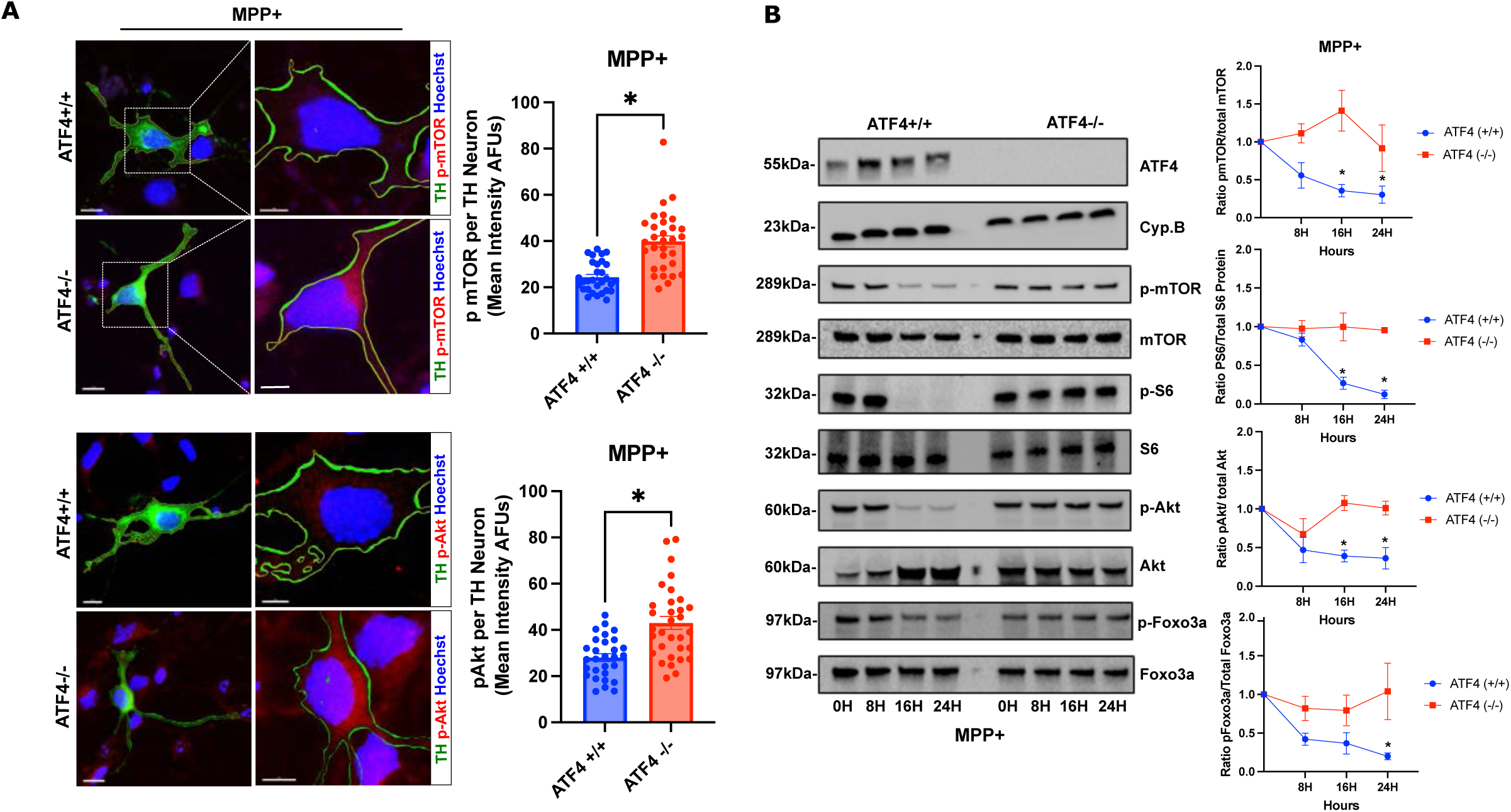
ATF4 regulates MPP+ mediated suppression of mTORC1/mTORC2 activity. A. *ATF4+/+* and *ATF4-/-* mesencephalic neurons treated with MPP+ (25μM) were fixed after 24 hours and co-immunostained for tyrosine hydroxylase (TH) and either p-mTOR (S2448) or p-AKT (S473) (scale bar= 15uM). Images on the right have been magnified and the outline of the TH-stained area highlighted to enable visualization of p-mTOR/p-AKT (red) fluorescence (scale bar=5uM). Confocal z-stacked images were acquired and the fluorescence intensity of p-mTOR (S2448) and p-AKT (S473) was quantified in dopaminergic (TH+) neurons from 3-independent experiments (n=29-32 cell per group, t-test, *p<0.05). **B.** *ATF4+/+* an *ATF4-/-* cortical neurons were treated with MPP+ (50μM) and protein was harvested at the indicated times and the levels of p-mTOR (S2448), p-S6 (S235/236), p-AKT (S473) and p-FOXO3a (S253) and corresponding total protein was assessed by western blot and quantified by densitometry. Quantification is reported as the ratio of phosphorylated protein/ total protein (n=5; 2way-ANOVA, *p<0.05).

**Supplemental Figure S2.**
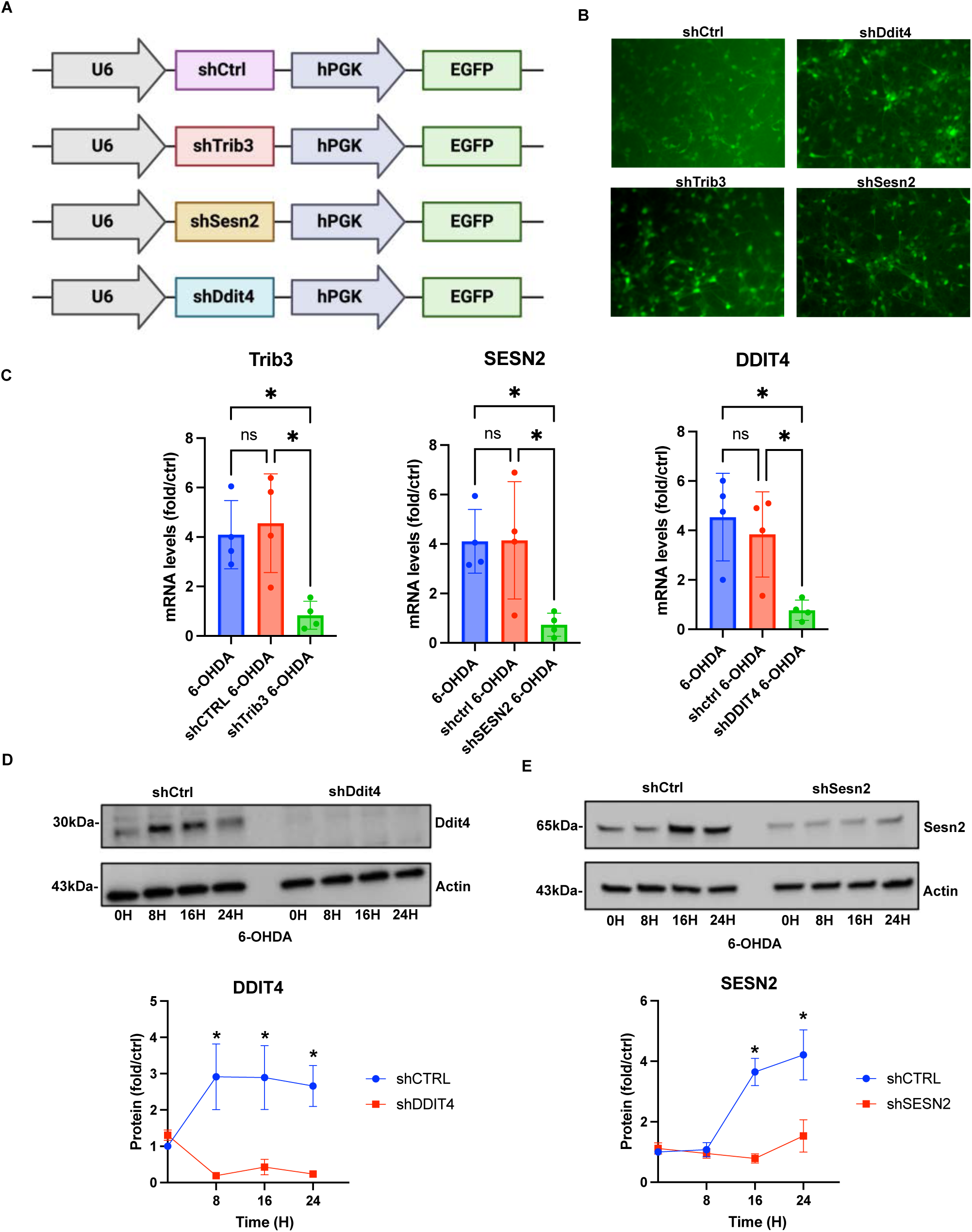
Validation of Lenti-shRNA vectors targeting SESN2, DDIT4 and Trib3. A. Schematic of Lenti-shRNA constructs co-expressing target specific shRNAs and EGFP from the U6 and hPGK promoters respectively. **B.** Representative image of cortical neurons transduced with Lenti-shRNA/GFP viruses for 5 days demonstrating high transduction efficiency (>90% GFP+). **C.** Cortical neurons were transduced with lenti-shRNAs targeting Sesn2, Trib3 or Ddit4 or a non-targeting lenti-shRNA control vector. Neurons were treated with 6-OHDA (10μM) for 16 hours and the mRNA levels of Sesn2, Trib3 and Ddit4 was determined by qRT-PCR. Data is reported as fold increase over untreated control cells from the same cultures (n=3; 1way- ANOVA *p<0.05). **D.** Cortical neurons were transduced with lenti-shRNA vector targeting Sesn2 or Ddit4 or a non- targeting lenti-sh-Ctrl vector. Neurons were treated with 6-OHDA (10μM) and SESN2 and DDIT4 protein levels were assessed at the indicated times by western blot and quantified by densitometry (n=3; 2-way ANOVA *p<0.05).

**Supplemental Figure S3.**
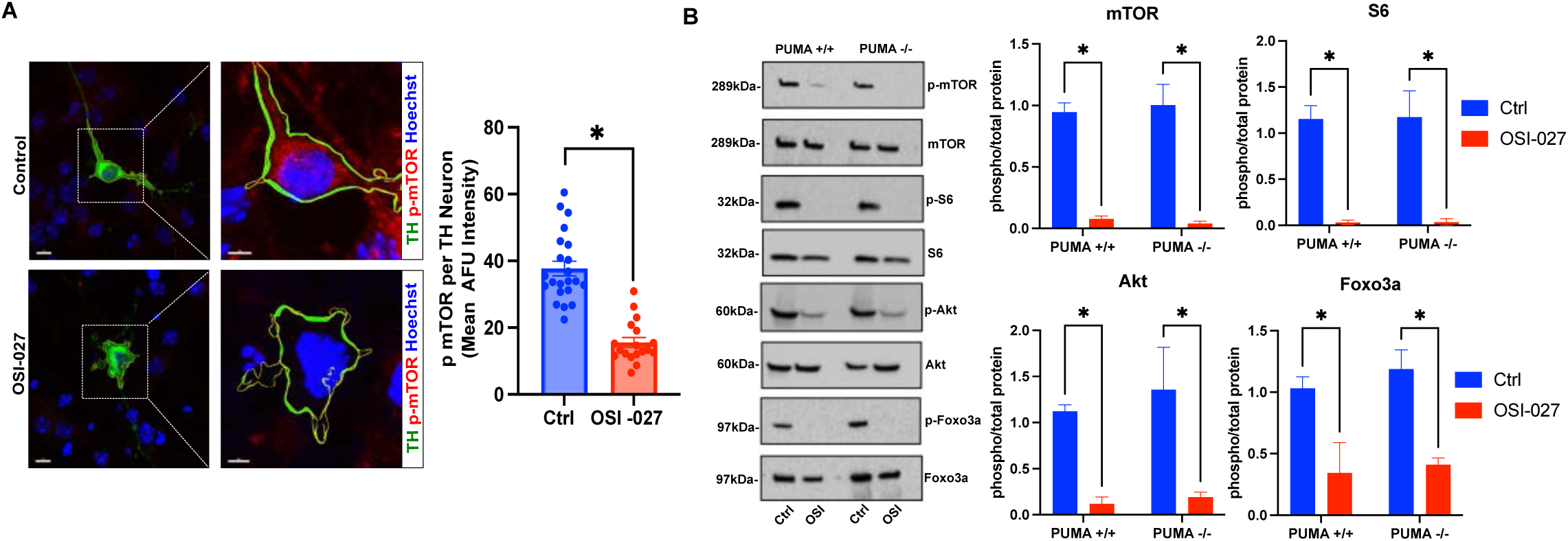
Validation of OSI-27 mediated suppression of mTORC1 and mTORC2 activity in neurons. A. Midbrain neurons were treated with the dual mTOR inhibitor OSI-027 (5μM) for 24 hours and then co-immunostained for p-mTOR (S2448) and tyrosine hydroxylase (left scale bar=15uM, right scale bar=5uM). Confocal images were acquired and the fluorescence intensity of p-mTOR (S2448) was quantified in dopaminergic (TH+) neurons from 3 independent experiments (n=18-22 cells per group; t-Test, *p<0.05). **B.** *PUMA+/+* and *PUMA-/-* cortical neurons were treated with OSI-027 (5μM) for 24 hours and the levels of p-mTOR (S2448), p-S6 (S235/236), p-AKT (S473), p-FOXO3a (S253) and corresponding total protein was assessed by western blot and quantified by densitometry. Quantification is reported as the ratio of phosphorylated protein/ total protein (n=3; 2way-ANOVA, *p<0.05).

**Supplemental Table 1.**
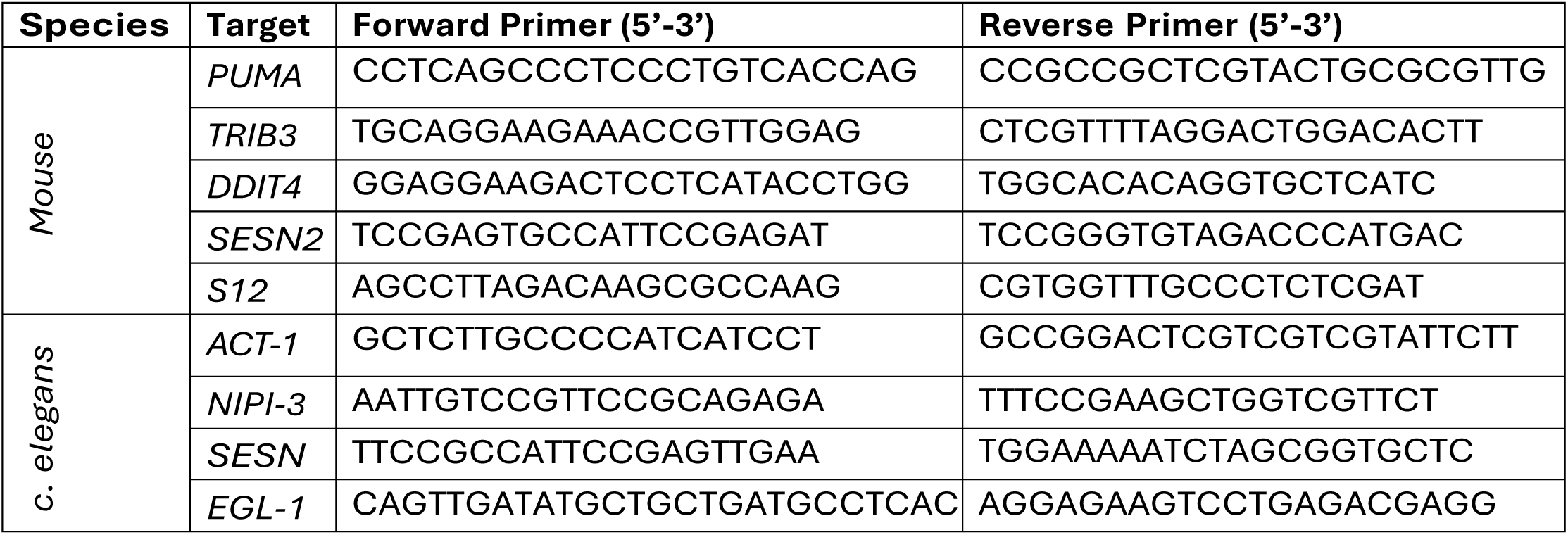
RT-qPCR Primer Sequences

**Supplemental Table 2.** Antibody information

| **Antibody** | **Dilution** | **Supplier** | **Catalog Number** |
| --- | --- | --- | --- |
| ATF4 | WB 1:1000 | Abcam | Ab184909 |
| Vinculin | WB 1:1000 | Biorad | MCA465GA |
| p-mTOR | WB 1:1000<br>IF 1:400 | Cell Signalling Technologies | **5536** |
| Total mTOR | WB: 1:1000 | Cell Signalling Technologies | 2972 |
| p-S6 | WB 1: 1000 | Cell Signalling Technologies | 4858 |
| Total S6 | WB 1: 1000 | Cell Signalling Technologies | 2317 |
| p-Akt | WB 1:1000<br>IF 1:400 | Cell Signalling Technologies | 9271 |
| Total Akt | WB 1:1000 | Cell Signalling Technologies | 9272 |
| p-Foxo3a | WB: 1000 | Cell Signalling Technologies | 2497 |
| Total Foxo3a | WB 1:2000 | ProteinTech | 66428-1-Ig |
| Tyrosine Hydroxylase (mouse) | IF 1:1000 | Millipore Sigma | MAB318 |
| Tyrosine Hydroxylase (rabbit) | IF 1:1000 | Millipore Sigma | AB152 |
| PUMA | WB 1:1000 | Cell Signalling Technologies | 98672 |
| Alpha-synuclein pSer129 | WB 1:1000<br>IF 1:500 | Abcam | Ab59264 |
| Sestrin2 | WB 1:1000 | ProteinTech | 10795-1-AP |
| DDIT4 | WB 1:1000 | Protein Tech | 10638-1-AP |
| Cyclophilin-B | WB 1:5000 | Abcam | Ab178397 |
| Alpha-tubulin | WB 1:2000 | Sigma | T9026 |
| Actin | WB 1:5000 | Sigma | A5441 |

